# Single-molecule evidence of Entropic Pulling by Hsp70 chaperones

**DOI:** 10.1101/2024.02.06.579217

**Authors:** Verena Rukes, Mathieu E. Rebeaud, Louis Perrin, Paolo De Los Rios, Chan Cao

**Affiliations:** Department of Inorganic and Analytical Chemistry, Chemistry and Biochemistry, University of Geneva; Geneva, 1211, Switzerland; Global Health Institute, School of Life Sciences, École Polytechnique Fédérale de Lausanne; Lausanne, 1015, Switzerland; Institute of Physics, School of Basic Sciences, École Polytechnique Fédérale de Lausanne; Lausanne, 1015, Switzerland; Institute of Bioengineering, School of Life Sciences, École Polytechnique Fédérale de Lausanne; Lausanne1015, Switzerland

## Abstract

Hsp70 chaperones are central components of the cellular network that ensure the structural quality of proteins. Despite their crucial roles in processes as diverse as the prevention of protein aggregation and protein translocation into organelles, their molecular mechanism of action has remained a hotly debated issue. Due to a lack of suitable methods, no experimental data has directly proven any of the models that have been proposed (Power Stroke, Brownian Ratchet, and Entropic Pulling). Recently, nanopores have emerged as a powerful tool to analyze the function of motor enzymes, as well as protein-protein interactions. Here, we used an *in vitro* single-molecule nanopore to mimic *in vivo* translocation of proteins, and to investigate the molecular mechanism of Hsp70. Our experiments demonstrate that Hsp70s forcefully extract polypeptide substrates that are trapped inside the pore. The forces they exert are strong at the molecular level, being equivalent to 46 pN over distances of 1 nm, and depend on the size of Hsp70. These findings provide unambiguous evidence supporting the Entropic Pulling mechanism of action of Hsp70s, thus solving a long-standing debate, and proposing a potentially universal principle governing diverse cellular processes. In addition, these results emphasize the utility of biological nanopores for studying protein function at the single-molecule level.

## Introduction

Molecular chaperones are essential components of the cell, as they protect organisms from the consequences of protein misfolding and/or aggregation by supervising the structural quality of proteins. The 70 kDa Heat Shock Protein (Hsp70) ATPase family is a central member of the chaperone repertoire, and is evolutionarily highly conserved (*1*). Its members perform a variety of cellular functions, including unfolding misfolded proteins, resolving protein aggregates, disassembling functional oligomers, and translocating proteins into mitochondria and the *Endoplasmic reticulum* (ER) (*2*, *3*). Because of their broad range of roles, Hsp70s are also implicated in many diseases and are consequently being investigated as drug targets for treatment, for instance in Parkinson disease and certain cancers (*4–6*).

Although Hsp70s are involved in a wide range of processes, the physical mechanism of their function remains uncertain. Protein translocation into organelles, for its geometric simplicity, is a well-suited setting to address this question. Proteins, synthesized in the cytoplasm, must pass through membrane channels to enter organelles, and often need to unfold from compact structures. This process is not spontaneous and requires assistance, which is provided by mitochondria- and ER-resident Hsp70s (*7*).

Two competing models for the molecular role of Hsp70s have originally been proposed: the *Power Stroke* (*8*) and the *Brownian Ratchet* (*9*). The Power Stroke model posits that Hsp70s bind to the translocating protein inside the organelle. Then, following ATP hydrolysis, they would undergo a conformational change resulting in an inward-directed pulling force on the Hsp70-bound substrate (Fig. S1a). The Brownian Ratchet (Fig. S1b), instead, suggests an entirely passive process: as a polypeptide segment emerges from the pore due to thermal fluctuations, binding by Hsp70 forbids any retro-translocation. Thus, only inward-directed thermal fluctuations are possible. The repetition of this process would progressively lead to protein import. Neither model is fully compatible with the current understanding of Hsp70s. For instance, the Power Stroke is in conflict with the ADP-bound structure of Hsp70, where the Substrate Binding Domain (SBD) and the Nucleotide Binding Domain (NBD) are connected by a very flexible linker that is unable to support a significant pulling force (*10*). Meanwhile, the Brownian Ratchet cannot explain the import forces observed in mitochondrial protein import (*11*) and is not easily applicable to processes with only one chaperone binding site, such as clathrin cage disassembly (*12*) and alpha-synuclein fibril disaggregation (*13*).

A third mechanism, *Entropic Pulling* (EP) (*14*, *15*), offers a more comprehensive physical explanation of the role of Hsp70 in protein translocation. EP suggests that the binding of Hsp70 to a protein segment emerging from the channel reduces the system’s entropy, because of the excluded volume between the chaperone and the close-by channel and membrane. Consequently, the bound Hsp70 attempts to increase its entropy, according to the second principle of thermodynamics, by moving away from the channel (Fig. S1c). This results in a force of entropic origin, pulling the bound protein with it. EP effectively combines aspects of the Power Stroke and Brownian Ratchet models, avoiding their limitations, and agrees with an increasing number of functional observations regarding Hsp70.

Nonetheless, there is no direct molecular proof of EP, nor experimental estimation of its strength. Previously, efforts to explore the molecular mechanism of Hsp70s experimentally have been made by observing the translocation of proteins into proteoliposomes (*9*) or mitochondria (*16*). In these bulk experiments, it was challenging to unambiguously select one model over the others, suggesting that more precise techniques are necessary. Single-molecule detection methods have recently opened new possibilities to investigate the functions of Hsp70 proteins (*17–20*). Among them, nanopores have emerged as powerful single-molecule tools. Besides their commercial use in DNA/RNA sequencing, they have also been demonstrated for characterizing protein interactions and functions (*21*, *22*). Motor enzymes with nucleic-acid substrates have been characterized at ultra-high spatiotemporal resolution, using single-molecule picometer resolution nanopore tweezers (*23*). In addition to its high resolution, nanopore technology has the advantage of being compatible with physiological conditions and does not require additional labeling of the investigated protein.

Here we have leveraged the similarities between the nanopore setup and *in vivo* translocation channels to study the physical mechanism of Hsp70s at the single-molecule level (Fig. 1). Like these *in vivo* channels, biological nanopore sensors are trans-membrane proteins. By incorporating them into artificial lipid membranes, a voltage can be applied across these pores, resulting in measurable ionic currents (*24*). In our system, we additionally used the applied potential to generate forces, against which the chaperone pulls a substrate protein. This mimics the presence of folded domains on the cytosolic side, that oppose the pulling of organellar Hsp70s during protein translocation. Also, *in vivo* translocation pores are characterized by the presence of abutting J-domain proteins (*25–28*). The J-domains of these proteins are essential to accelerate the chaperone ATPase cycle, resulting in the locking of Hsp70 onto the client (*29*). We reproduced this setup by devising a protein construct with an N-terminal J-domain, to recruit Hsp70, and a C-terminal negatively charged polypeptide that is trapped inside the pore by the applied voltage.

**Fig. 1.**
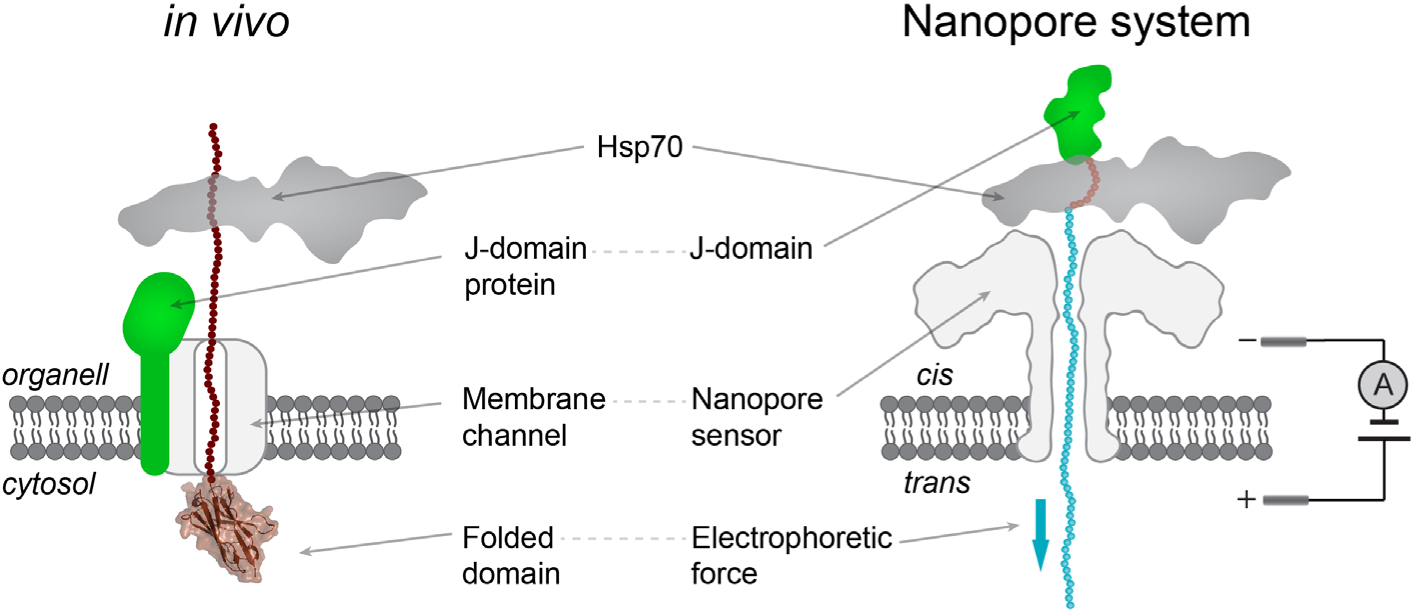
Illustration of the designed system compared to an example of *in vivo* protein transport mediated by Hsp70s. The nanopore functions as the membrane channel (light grey). Instead of a folded domain as *in vivo* (brown), the enzyme is working against a voltage-induced force in our system that acts on the charged part of the designed substrate (blue).

## Results

### Monitoring Hsp70-substrate binding in nanopore experiments

In our designed system we monitored the escape of different substrates from the pore, by measuring the time needed for escape, which was greatly facilitated by the binding of Hsp70 (here the bacterial homolog DnaK). Escapes also happened faster for constructs of increasing size. This is in agreement with the EP predictions that larger molecules induce a greater reduction of the entropy of the system, and thus larger forces. Using theoretical modeling, we estimate that binding of Hsp70 can lower the energy barrier for escape by about 10 *k*_B_*T* (6 kcal/mol). These results cannot be explained by the Power Stroke or the Brownian Ratchet, thus establishing EP as the molecular mechanism governing the action of Hsp70s.

To mimic Hsp70-mediated pulling *in vitro*, we designed the protein construct Jdp5Nt (Fig. 2a), which functions as a substrate for DnaK (Fig. 2b). Jdp5Nt consists of three parts: (i) the folded J-domain (amino acids 1-70) of bacterial DnaJ; (ii) a well-established DnaK binding peptide, the p5 sequence, ALLLSAPR (*30*); (iii) a non-structured tail of 60 amino acids with a evenly distributed charge of -20 at pH 7.4 (Fig. S2). This tail allows the electrophoretic capture of Jdp5Nt into the nanopore, with the consequent reduction of the ionic current (Fig. 2c). The N-terminal J-domain sterically prevents complete translocation, while simultaneously mediating substrate binding by DnaK. The proper functionality of DnaK was confirmed through luciferase unfolding assays (Fig. S3).

**Fig. 2.**
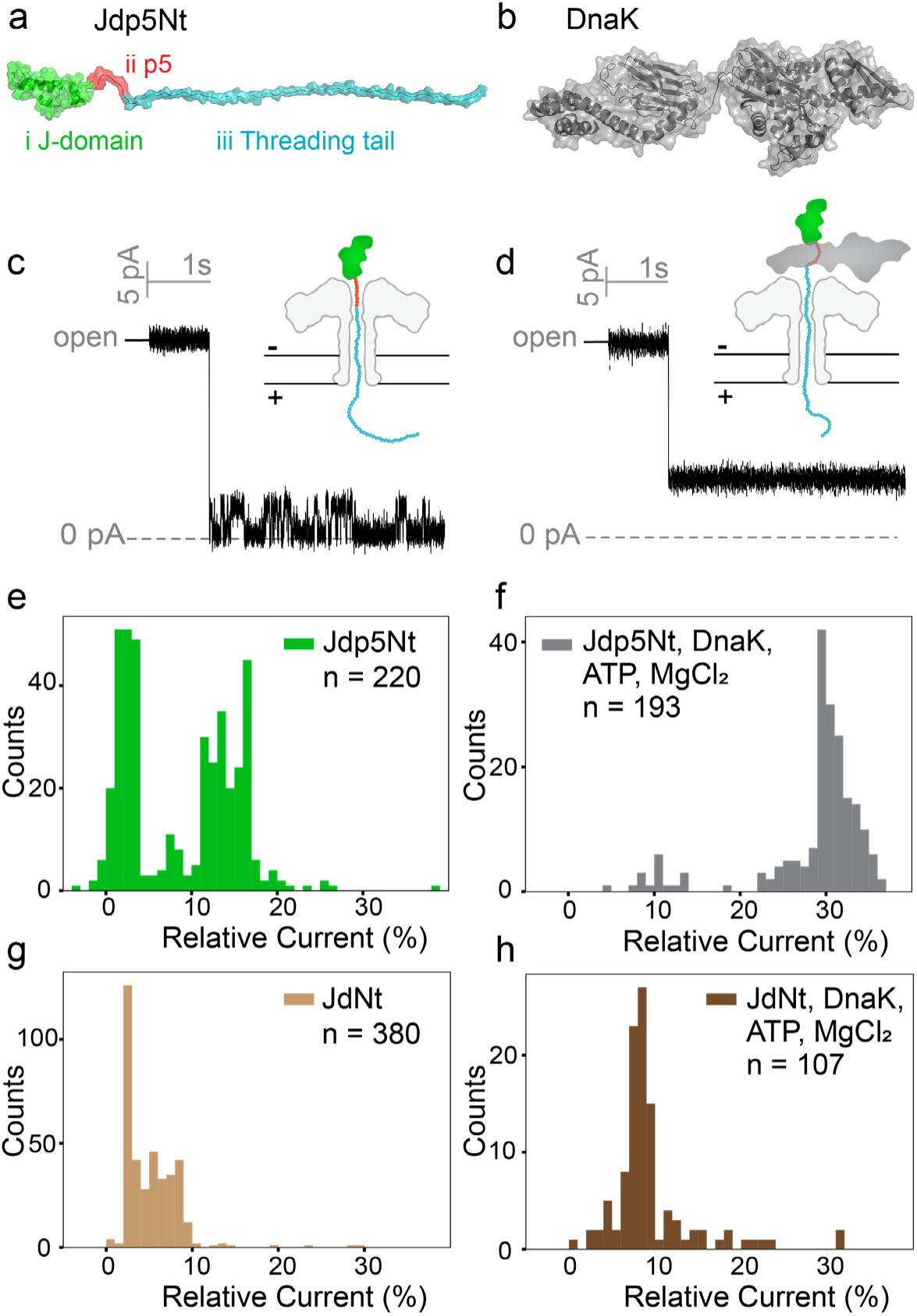
Monitoring Hsp70-substrate binding in nanopore experiments. (a) Atomistic representation of the Jdp5Nt construct. (b) Structure of DnaK in its ADP-bound state (PDB 2KHO). (c) Representative current signal stemming from measurements of Jdp5Nt. During experiments, Jdp5Nt is captured and trapped inside the nanopore as illustrated. (d) Representative current signal in the presence of Jdp5Nt, DnaK and ATP (complex-formation-condition). Distribution of the gaussian fitted values for experiments with only Jdp5Nt present (e), in the complex condition (f), with only JdNt present (g) and with JdNt, DnaK and ATP (h). The current of each trapping event was fitted with two Gaussians for Jdp5Nt or one Gaussian in all other conditions.

Through analysis of the ionic current signatures and escape voltages we confirmed that the complex of Jdp5Nt and DnaK can be trapped and measured by the nanopore sensor (as illustrated in Fig. 2d). We found that analyte-trapping in the pore produced characteristic current signals that can be used to identify the captured analyte. Without the chaperone, we observed a bimodal current with one state at 3% ± 3% (relative to the open-pore current, corresponding to 100%) and another at 14% ± 4% (Fig. 2e; see Methods for the determination of the currents). In the presence of DnaK and ATP, the current distribution peaked at 29% ± 7% (Fig. 2f). In some instances, we observed a transition from these higher currents to the ones typical for Jdp5Nt alone (Fig. S4). This is in agreement with the expected scenario of DnaK occasionally disengaging from Jdp5Nt. To control the role of the p5 peptide for DnaK binding, we also measured the current distribution for the JdNt construct, which lacks the p5 peptide. This substrate alone yielded a one-state current signal of 5% ± 3% (Fig. 2g). As expected from the absence of p5, currents for JdNt with DnaK and ATP (9% ± 5%, Fig. 2h), showed no significant shift compared to JdNt alone. To further confirm that Jdp5Nt engages in complex with DnaK via its p5 binding site, escape voltage experiments were carried out (Note S1, Fig. S5). Addition of DnaK and ATP facilitated the escape against higher voltages of Jdp5Nt, which is not the case for JdNt.

### Experimental evidence of entropic pulling by Hsp70

When charged polymers are trapped in the pore by an applied voltage, they can escape back to the *cis* compartment, against the trapping force. It has been previously determined that the average escape time depends exponentially on the applied voltage, a hallmark of thermally-activated processes over an energy barrier that is in turn proportional to the voltage (*31*). In keeping with these observations, the average escape time of Jdp5Nt increased roughly exponentially for increasing voltages (Fig. 3a; see Methods and Fig. S6 for details of the measurement). JdNt showed similar escape times to Jdp5Nt (2.2 s and 2.4 s, respectively, at 5 mV Table S1) confirming that the p5 sequence had no relevant influence on the escape. In the presence of DnaK and ATP, the escape times of Jdp5Nt were reduced by orders of magnitude even against higher applied voltages. Nonetheless, they still increased exponentially with the applied voltage (Fig. 3a). Thus, the binding of DnaK to the Jdp5Nt polypeptide resulted in a reduction of the energy barrier that hinders escape. Additionally, we designed the construct NBDNt, which consists of the nucleotide binding domain of DnaK with the threading tail of Jdp5Nt added at the C-terminal (Fig. S2 and Methods). The escape rates of NBDNt were intermediate to the ones of Jdp5Nt with and without DnaK and ATP, for intermediate applied voltages (Fig. 3a). The compared systems are illustrated in Fig. 3b-d.

**Fig. 3.**
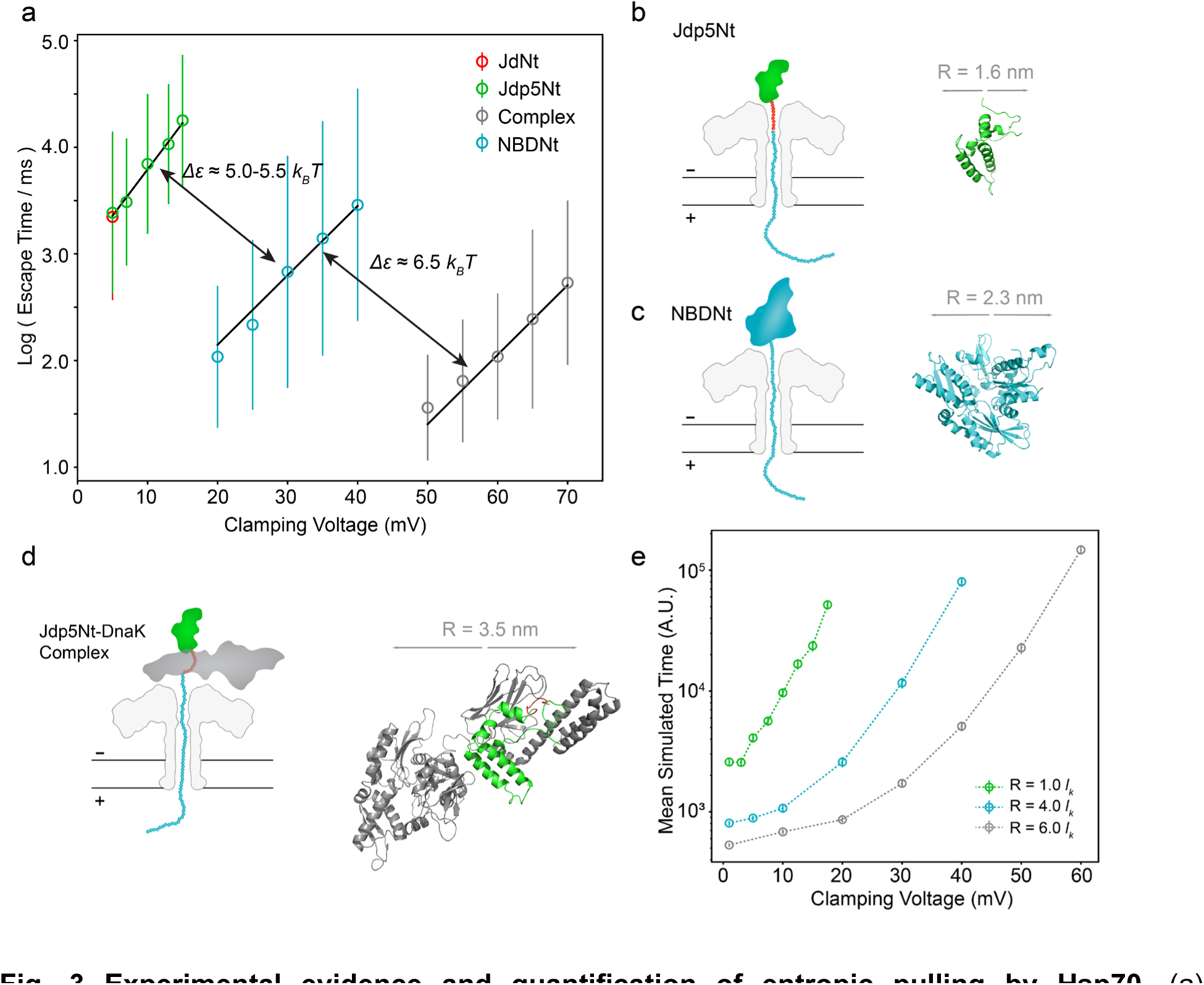
Experimental evidence and quantification of entropic pulling by Hsp70. (a) Logarithmized escape times of the systems (measurement details in methods and Fig. S6). The log values of each system at various clamping voltages were averaged, standard deviation is indicated. The mean values in ms were fitted to Eq.2 (black lines, table S2). (b)-(d) illustrations of the measured systems Jdp5Nt, NBDNt and Complex, respectively. Snapshots from MD simulation of each folded domain are shown with the calculated radii. (e) Simulated escape times (in arbitraty units, A.U.) using different sizes of the domain above the pore (in units of the Kuhn length of unfolded polypeptides, lK). Details of the model in Note S2.

The exponential dependence of the average escape-time 𝜏𝜏̅ on the applied voltage can be modeled using Kramer’s theory for thermally-activated escape over an energy barrier (*32*), with

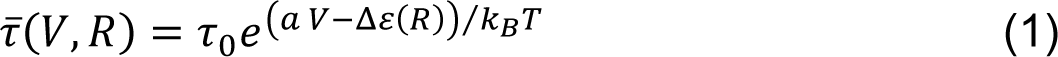

where 𝜏_0_ captures various intrinsic properties of the pore, of the trapped polypeptide and of their interactions, 𝑎 𝑉 is the linear increase of the energy of the barrier due to the applied voltage *V*, Δ𝜀(𝑅) is the reduction of the barrier energy due to the presence of a bulky N-terminal domain of radius *R*, k_B_ is the Boltzmann constant and *T* the absolute temperature. The parameter 𝑎 describes how the pore, the trapped polypeptide, and their physical and chemical features determine the electrophoretic force generated by the applied voltage. In Fig. 3a the experimental data were fitted to the exponential formula

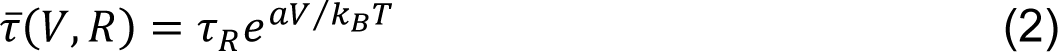

where 𝜏_𝑅_ = 𝜏_0_𝑒^−Δ𝜀(𝑅)/𝑘𝐵𝑇^. The corresponding values of 𝑎 and 𝜏_𝑅_ are reported in Table S2. Everything else being equal, these results support the entropic pulling mechanism, which posits that increasing the size of the N-terminal domain results in stronger pulling forces that reduce the escape times (*14*).

### Quantification of Entropic Pulling

We quantified the effect of construct size on the escape time, by estimating the energy barrier for escape, Δ𝜀(𝑅), in each system. Using Eq.1 and taking the ratio of the average escape times using the fitted values at a fixed voltage but for different constructs (Fig. 3a), we obtained Δ𝜀(𝑅_𝐷𝑎𝐷_) − Δ𝜀(𝑅_𝑁𝐷_) ≈ 6.5 *k*_B_ *T* (3.9 kcal/mol = 26.2 pN nm) and Δ𝜀(𝑅_𝑁𝐷_) − Δ𝜀(𝑅_𝐽−domain_) ≈ 5.0-5.5 *k*_B_ *T* (3.0-3.3 kcal/mol = 20.0-22.0 pN nm. Because fitting of Jdp5Nt and NBDNt data resulted in different values for 𝑎, a range is obtained for Δ𝜀(𝑅_𝑁𝐷_) − Δ𝜀(𝑅_𝐽−domain_). It represents the estimates for the smallest and largest measured values for Jdp5Nt. Thus, the NBD of DnaK reduced the barrier by about 5.0 *k*_B_*T* (taking a lower, conservative value) from the value of Jdp5Nt, and the complex with DnaK reduced it by a further 6.5 *k*_B_*T* from the barrier value in the presence of NBDNt. In total, binding of DnaK reduced the energy barrier by roughly 11.5 *k*_B_*T* (6.9 kcal/mol = 46 pN nm).

It was previously predicted, that the energy contribution of the bulky domain during EP is proportional to the square of its radius (*14*). We found that, relative to Jdp5Nt, the DnaK complex reduced the energy barrier 2.3 times more than the NBD. As a consequence, our calculations predict that the radius of the DnaK complex should be roughly √2.3 ≈ 1.5 times larger than the one of the NBD alone. We thus performed molecular dynamics simulations of the J-domain, NBD and of a full-length ADP-bound DnaK to estimate their radii (Methods, Fig. S7). Snapshots of each simulated system are shown in Fig. 3b-d. In good agreement with our predictions, the average radius of the simulated DnaK in complex with Jd was 1.5 times larger than the one of NBD-ADP (Table S3).

While our data were generally fitted well by an exponential, some deviation was particularly apparent in the case of the DnaK complex at the lowest measured clamping voltages (Fig. 3a). This behavior is predicted for small, negligible energy barriers (<< *k_B_T*), when escape times are dominated by diffusion in the pore, thus approaching a constant value (*33*). Without attempting to capture all details of the polypeptide-pore interactions, nor the exact shape of the electrophoretic trapping potential, we were able to model the behavior of our experimental system (see Note S2 and Fig. S8 for details of the model). We simulated the escape process for different voltages and sizes of the N-terminal domain. The results reproduced the behavior of the escape time we found in nanopore experiments (Fig. 3e), confirming that the binding of DnaK reduced the energy barrier enough to make it smaller than *k_B_T* at low applied voltages.

## Discussion

Although the Hsp70 family of ATP-dependent chaperones plays crucial roles in extremely diverse biological processes in all cells, the underlying mechanism of their functions has so far not been identified unambiguously. In this study, we have developed a nanopore-based system to simulate *in vivo* protein transport mediated by Hsp70s, and monitor this process *in vitro* at the single-molecule level. We found that binding of Hsp70 greatly facilitated the escape of a polypeptide from the pore. By comparing its effects with domains of different sizes, we estimated that the binding of DnaK lowers the energy barriers that must be overcome to escape from the pore by as much as 11-12 *k*_B_*T*. In the context of protein import this translates in the acceleration of the unfolding of folded domains on the cytoplasmic side of the organellar membrane, as proposed originally (*14*), and can explain the observation that folded and unfolded domains are imported at the same rate (*11*).

Out of the proposed mechanisms, Power Stroke, Brownian ratchet, and EP, our experiments confirm EP as the molecular mechanism for force generation by Hsp70. Our findings are not compatible with the Power Stroke model because the effect is also present for the passive construct NBDNt in the presence of ADP, which is not hydrolysable, and does not undergo any specific lever-arm conformational transition. The Brownian Ratchet also fails to explain our results, since even the smallest measured constructs, Jdp5Nt and JdNt, forbid their full translocation and no further size-dependent effects are predicted by the Brownian Ratchet. Furthermore, our system does not allow the progressive binding of Hsp70 molecules, because there is a single binding site on Jdp5Nt (p5 peptide) or, in the case of NBDNt, there are no molecules in solution that could bind the construct. This direct confirmation of EP and its quantitative assessment reveal the key principle that underlies the function of Hsp70s molecular chaperons, and other biological processes (*34*). This work also highlights the versatility of nanopore single-molecule method to study and quantify the interactions and mechanical properties of complex biological molecules, which enables measurements that are currently beyond the reach of other tools.

## Acknowledgments

This work was supported in part using the resources and services of the Protein Production and Structure Core Facility at the School of Life Sciences of EPFL. The authors also thank K Lau for support and scientific discussion concerning protein production.

## Author contributions

C.C., P.D.L.R., M.R. and V.R. conceived the project and designed the study. V.R. performed all experiments and signal processing. P.D.L.R. and V.R. did data analysis.

P.D.L.R. build the physical model and performed simulations. M.R. carried out luciferase unfolding assays. L.P. performed molecular dynamics simulations. V.R., C.C., and P.D.L.R. interpreted the data. V.R., C.C., and P.D.L.R. wrote the manuscript with input from all authors.

## Data and materials availability

All data are available in the main text or the supplementary materials.

## Materials and Methods

### Protein production

DnaK, Jdp5Nt, JdNt and NBDNt were produced by the Protein Production and Structure Core Facility at École polytechnique fédérale de Lausanne. Sequences for Jdp5Nt, JdNt and NBDNt were cloned in pEt-29b(+) vectors by GenScript custom gene service. The plasmid for DnaK production was provided by Dr. Mathieu Rebeaud. Jdp5Nt and JdNt were produced in *E. coli* BL21(DE3) pLysS. DnaK and NBDNt were produced in *E. coli* BL21(DE3). Jdp5Nt, JdNt and NBDNt were purified by His-tag chromatography and a subsequent size exclusion chromatography. DnaK was purified through SUMO-tag purification with subsequent tag-cleavage with ULP1 and size exclusion chromatography. After several washes with high salt buffer (+150 mM KCl, 20mM Imidazole and 5 mM ATP), N-terminal His10-SUMO (small ubiquitin-related modifier) Smt3 tag was cleaved with Ulp1 protease (2mg/ml, 300 μl, added to beads with buffer (20 mM Tris-HCl pH 7.5, 150mM KCl, 10mM MgCl2, 5% glycerol). Digestion of His10 Smt3 was performed on the Ni-NTA resin by, His6-Ulp1 protease. After overnight digestion at 4°C, the unbound fraction was collected (which contains only the native proteins). Pure fractions were pooled, concentrated by ultrafiltration, aliquoted in appropriate buffer, and stored at -80 °C.

### Nanopore measurements

Nanopore measurements were carried out in 300 mM KCl, 10 mM HEPES, pH 7.4 (except for initial escape voltage measurements in 1 M KCl, 10 mM HEPES, pH 7.4). Lipid bilayers of 1,2-diphytanoyl-sn-glycero-3-phosphocholine (DPhPC) were formed on the 50 µm apertures of MECA 4 Recording Chips (Ionera, Germany) and small amounts of activated aerolysin pore were added to the *cis* chamber to allow single pore insertion. Data was acquired at 50 kHz with an amplifier gain that allowed recordings between -200 pA and 200 pA using an Orbit mini device (nanion technologies, Germany). Temperature control of the instrument was set to 25°C for all measurements. Each experiment was carried out several times and with >5 individual pores. Jdp5Nt and JdNt were added to 2 µM final concentration when measured individually. For complex experiments, most experiments contained 800 nM DnaK, 200 nM Jd(p5)Nt, 1 mM ATP and 1 mM MgCl_2_. NBDNt was measured at concentrations of 0.5 or 1 µM. For NBDNt-ADP/NBDNt-ATP measurements, the buffer contained 1 mM ADP/ATP and 10 mM MgCl_2_.

To determine against which approximate voltages a given analyte is likely to escape the aerolysin pore, analytes were first trapped inside the pore by applying a voltage ≥ 100 mV. The trapping resulted in an instantaneous blockage of the current through the pore. Subsequently, the applied voltage was lowered in 10 mV increments and each voltage was held for 2 s. When the analyte escaped from the pore, the measured current jumped back to the open pore current at the given escape voltage.

For all measurements of escape time, the analytes were captured at 140 mV (Fig. S6). After trapping for at least 4 s the voltage was shifted to the clamping voltage. The time needed for an analyte to escape against the clamping voltage was subsequently determined by hand in ClampFit. In the case of clamping voltages <20 mV data was gaussian-filtered to 50-100 Hz resulting in a detection limit of ∼50 ms. For higher clamping voltages filtering to 200-500 Hz was sufficient to determine transitions in current states which allowed to determine escapes as short as ca. 5 ms.

### Data analysis

To investigate the residual current during the trapping of different analytes data was first gaussian filtered in ClampFit to 500 Hz. A custom python script was subsequently used to extract the data points of 3 s of when an analyte was held in the pore with 140 mV voltage bias. Histograms of these data points were plotted and gaussian fitted for the individual trapping events. The current of each event was fitted with two Gaussians for JdptNt or one Gaussian in all other conditions. Escape voltage and escape time values were extracted by hand using ClampFit software. The values of the escape times of each analyte and clamping voltage were averaged.

### Luciferase refolding assay

Luciferase activity was measured as described previously(*1*, *2*). In the presence of oxygen, luciferase catalyzes the conversion of D-luciferin and ATP into oxyluciferin, CO_2_, AMP, PPi, and hν. Generated photons were counted with a Victor Light 1420 Luminescence Counter from Perkin-Elmer (Turku, Finland) in a 96-well microtiter plate format.

### Molecular dynamics simulations

The J-domain alone (structure predicted with alpha fold(*3*, *4*), no p5, no threading tail), the complex of Jdp5 (structure predicted with alpha fold, no threading tail) and DnaK (https://doi.org/10.2210/pdb2kho/pdb), and NBD in the ADP bound state (https://doi.org/10.2210/pdb3ay9/pdb) were simulated. All systems were built with the CHARMM-GUI (*5*, *6*), solution builder. Where necessary, purification tags were added in Pymol (sequence: MKHHHHHHHHHHGSAN). On every system, restrains were applied during equilibration: for the backbone heavy atoms, 400 kJ mol^−1^ nm^−2^, for the side chain heavy atoms, 40 kJ mol^−1^ nm^−2^.

Each of the systems was put in a periodic box of water containing 0.3 M of KCl. Minimization was run to get the maximal force value below 1000 kJ mol^−1^ nm^−1^. Each of the systems was equilibrated for 125000 steps with a time-step of 1 fs before production. No particular restrains were applied during production. Root mean square deviation and radius of gyration were calculated using GROMACS(*7*) software and averaged over frames.

## Supplementary Text

### Note S1 Measurements of escape voltage

Measurements of escape voltages allow us to assess, against which voltage biases each analyte is likely to escape the nanopore. As illustrated in Figure S5, the voltage is decreased in 10 mV steps during trapping events until the analyte exits the pore towards *cis* at the escape voltage. Jdp5Nt or JdNt on their own escape most prevalently at voltages between 0 mV and 30 mV. Long duration blockages from DnaK end at more broadly distributed voltages. This might be because the designed constructs are trapped via their negatively charged tail while capture and escape of DnaK is less distinctive. In the presence of Jdp5Nt, DnaK, MgCl_2_ and ATP the distribution of escape voltages is again more concentrated peaking at 80 mV. Escapes in the lower voltage regime below 50 mV are rare. If, however, JdNt is measured together with DnaK, MgCl_2_ and ATP the distribution of escape voltages resembles a superposition of the ones of JdNt and DnaK with no indication of a complex being measured. This further identifies the p5 sequence as the binding site of a DnaK-Jdp5Nt complex and shows that the escape behavior of this complex differs from that of Jdp5Nt. In the presence of the chaperone, Jdp5Nt can escape the aerolysin nanopore against a higher voltage bias.

Escape voltages were also measured with NBDNt. Surprisingly, in its apo-state NBDNt escaped mostly at negative voltages, requiring even lower voltages than Jdp5Nt. Since the NBD is known to hydrolyze ATP and efficiently bind ADP when isolated from the SBD(*8*), measurements were conducted in the presence of 1 mM of ADP or ATP together with 10 mM MgCl_2_. In both conditions the escape voltages shifted to higher positive values around 30-60 mV. This could be explained by the higher stability of lobe II of the NBD upon nucleotide binding(*9*). Since this lobe is located at the C-terminus, where the threading tail of NBDNt was added, the stability of this lobe is likely critical in nanopore experiments. We believe the unfolding of this lobe followed by partial unfolding of NBDNt in its apo-state is the cause of the low escape voltages of apo-NBDNt.

### Note S2 Simulations of the escape process

The geometry of the system has been chosen such that:

1) 𝐿 is the length of the polypeptide and 𝑙𝑙_𝑃𝑃_ is the length of the pore.
2) 𝑥 is the length of the polypeptide that emerges from the pore on the *cis* side.
3) 𝑥 = 𝐿: the polypeptide has completely escaped the pore.
4) 𝐿 − 𝑙_𝑃_ ≤ 𝑥 ≤ 𝐿 : the polypeptide has a part inserted in the pore but does not emerge on the *trans* side.
5) 0 ≤ 𝑥 ≤ 𝐿 − 𝑙_𝑃_: the polypeptide occupies the full length of the pore and emerges on the *cis* side by a length 𝑥, and on the *trans* side by an amount 𝐿 − 𝑙_𝑃_ − 𝑥.
6) 𝑙_𝐾_ is the Kuhn length of an unfolded polypeptide (about 2-3 amino-acids (*10*, *11*)).

The polypeptide moves in an energy landscape 𝐸(𝑥) that is the sum of three contributions (Fig.S8):

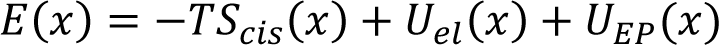

1) The length of the polypeptide in the *cis* side contributes by its entropy, which increases with the lenth as

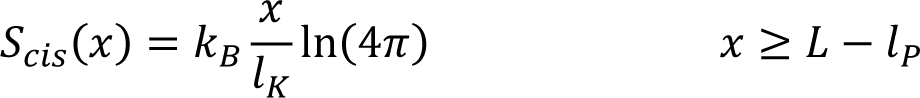 Here we have assumed that the polypeptide can be considered as a Freely Jointed Chain composed of a number of segments which is given by the length of the polypeptide emerging on the *cis* side, 𝑥, divided by the Kuhn length. We have also assumed that each segment can freely rotate, and that thus its entropy is just 𝑘_𝐵_ ln(4π). This contribution applies if the polypeptide emerges only from the *cis* side. As soon as it emerges also on the *trans* side, the amount of length lost on one side is gained on the other and the two effects mutually cancel out.
2) Considering the polypeptide to be uniformly charged with charge density 𝜆, in the presence of an applied voltage 𝑉 the electrostatic potential 𝑈(𝑥) is:

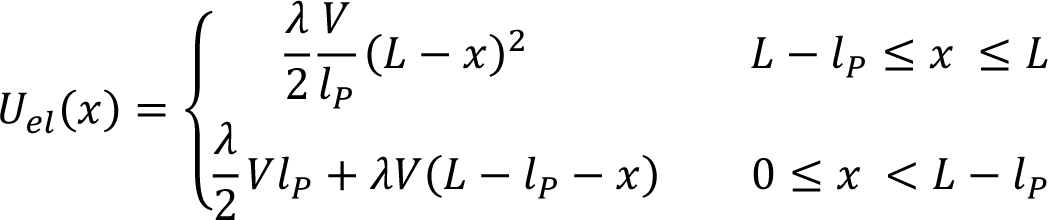
3) The entropic pulling contribution 𝑈_𝐸𝑃_(𝑥) is (*12*):

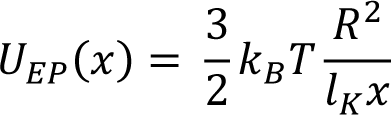

where 𝑅 is the radius of the bulky, N-terminal domain (or of the bound DnaK).

The overall parameters we have used to compute the potential were:

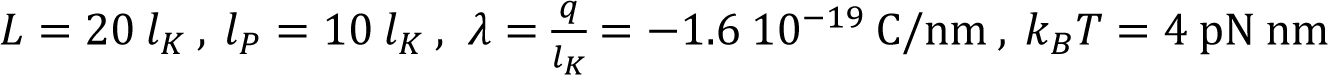

The energy 𝐸(𝑥) for 𝑉 = 55 mV and 𝑅 = 10 𝑙_𝐾_, as an example, is represented in Fig.S9.

The escape process was simulated as a random walk in the interval 0 ≤ 𝑥 ≤ 𝐿. The walker started at a position 𝑥_0_ and in a time-step it could move from 𝑥 of an amount Δ𝑥 to the left with probability

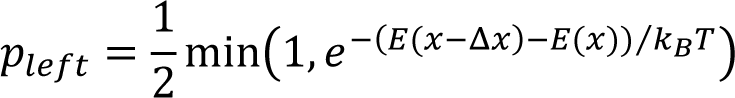

and to the right with the probability

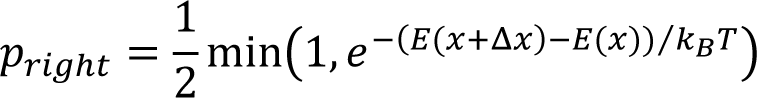

and would not move with probability *p_not move_* = 1 − *p_left_ p_right_*.

The random walk started from an initial random position 𝑥_0_ that was chosen by running the simulation with a high trapping voltage (140 mV) for 10’000 time-steps. The applied voltage was then lowered to the desired value, and the simulation was run until 𝑥 > 𝐿, and the escape time, namely the number of time-steps (in arbitrary units) was recorded. The average escape time was estimated over 100 runs of the simulation (𝑥_0_ was re-generated for each run) for each applied voltage and each value of the radius 𝑅.

---

**Fig. S1.**
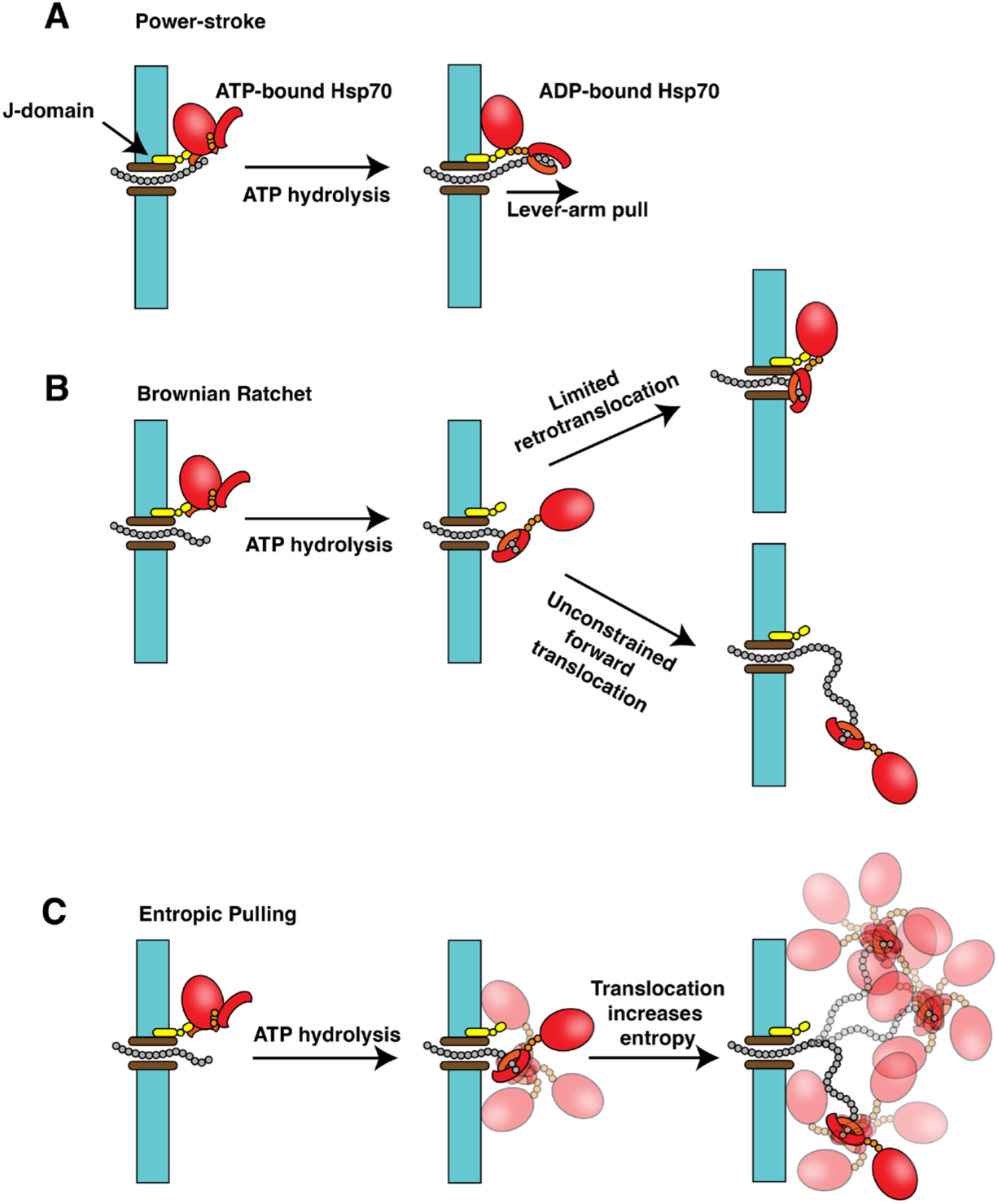
Proposed models for Hsp70s mechanism of function.

**Fig. S2.**
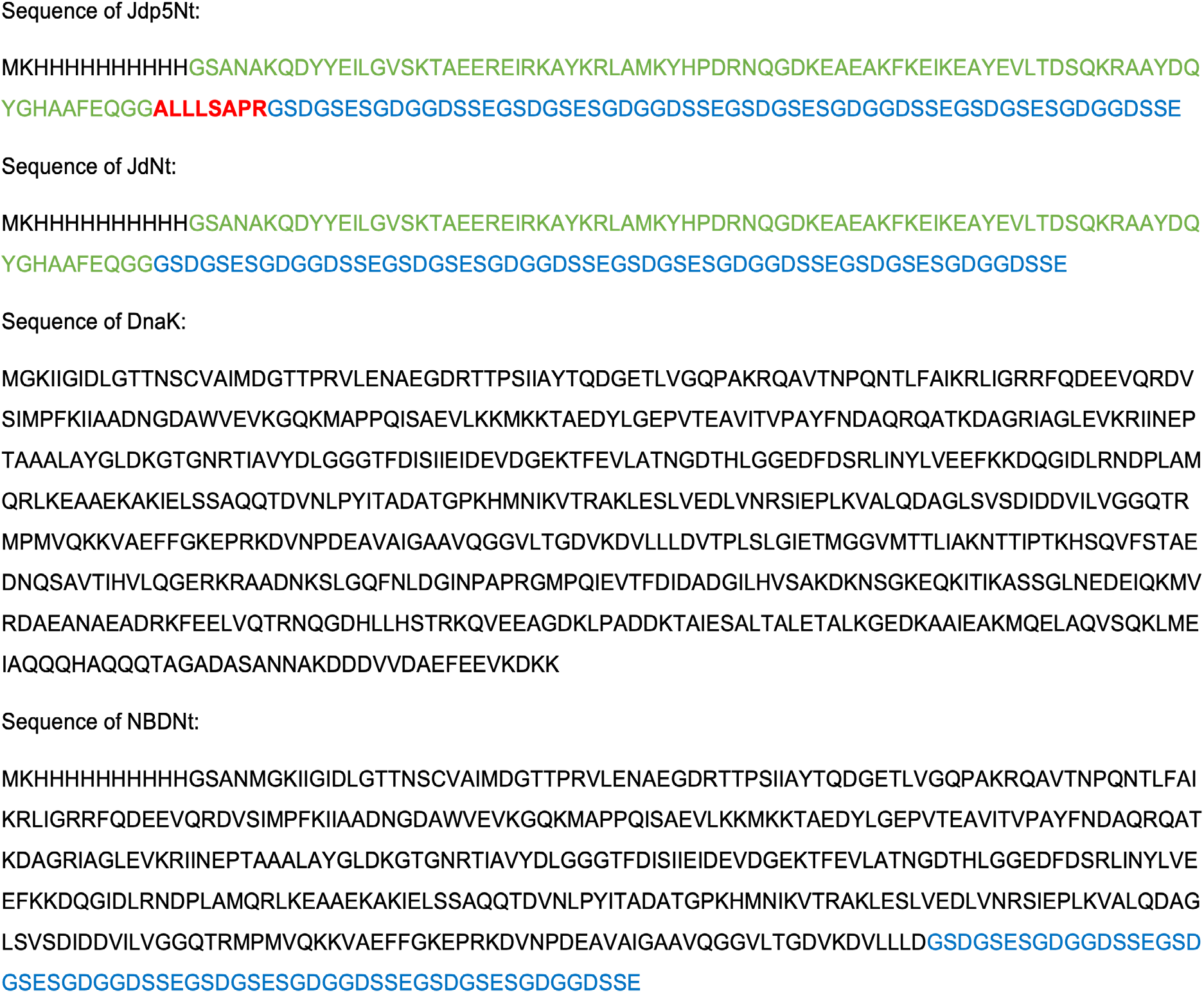
Amino acid sequences of the proteins used in experiments.

**Fig. S3.**
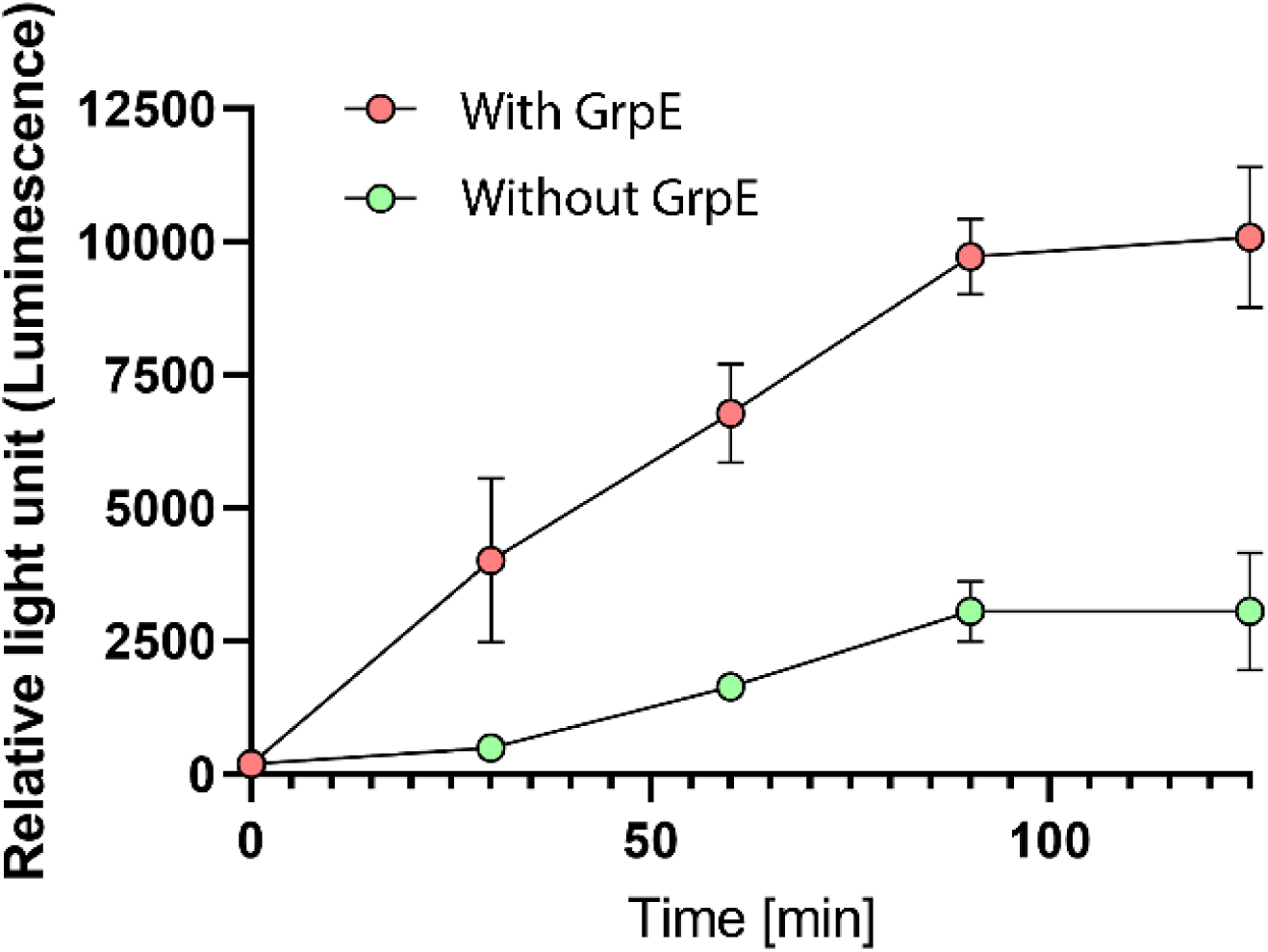
Time-dependent reactivation of urea-pre-denatured Luciferase that was incubated with (red) or without (green) GrpE. 50 µM Luciferase was first inactivated for 2 min in 6M Urea, then diluted 125 folds in buffer (25 mM HEPES-KOH pH 7.5, 150 mM KCl, 10 mM MgAc2, 1mM DTT, and 5 mM ATP) at 30°C without any chaperone for 30 minutes. The reaction mix was then diluted 2 folds (final Luciferase concentration: 200 nM) and supplemented with the fully active chaperone mix (4 µM DnaK, 1 µM DnaJ with or without GrpE), to reactivate the aggregated luciferase at 25 °C. The error bars represent the SD between replicates (n = 3).

**Fig. S4.**
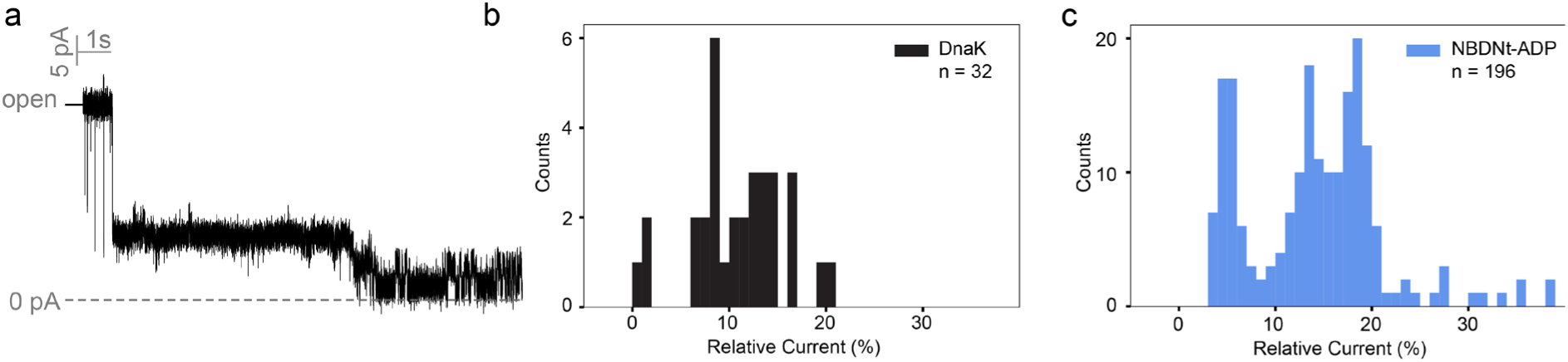
Current traces were acquired in 300mM KCl at 140 mV potential applied to the trans chamber filtered to 500 Hz. (a): example trace recorded in a condition with Jdp5Nt, DnaK, MgCl_2_ and ATP. After trapping a high current state associated with the Jdp5Nt-DnaK complex is observed, followed by the signal typical for Jdp5Nt. (b): Distribution of the fitted current values that were occasionally observed after addition of only DnaK. (c): Distribution of the fitted current values stemming from NBDNt in the presence of 1 mM ADP 10 mM MgCl_2_.

**Fig. S5.**
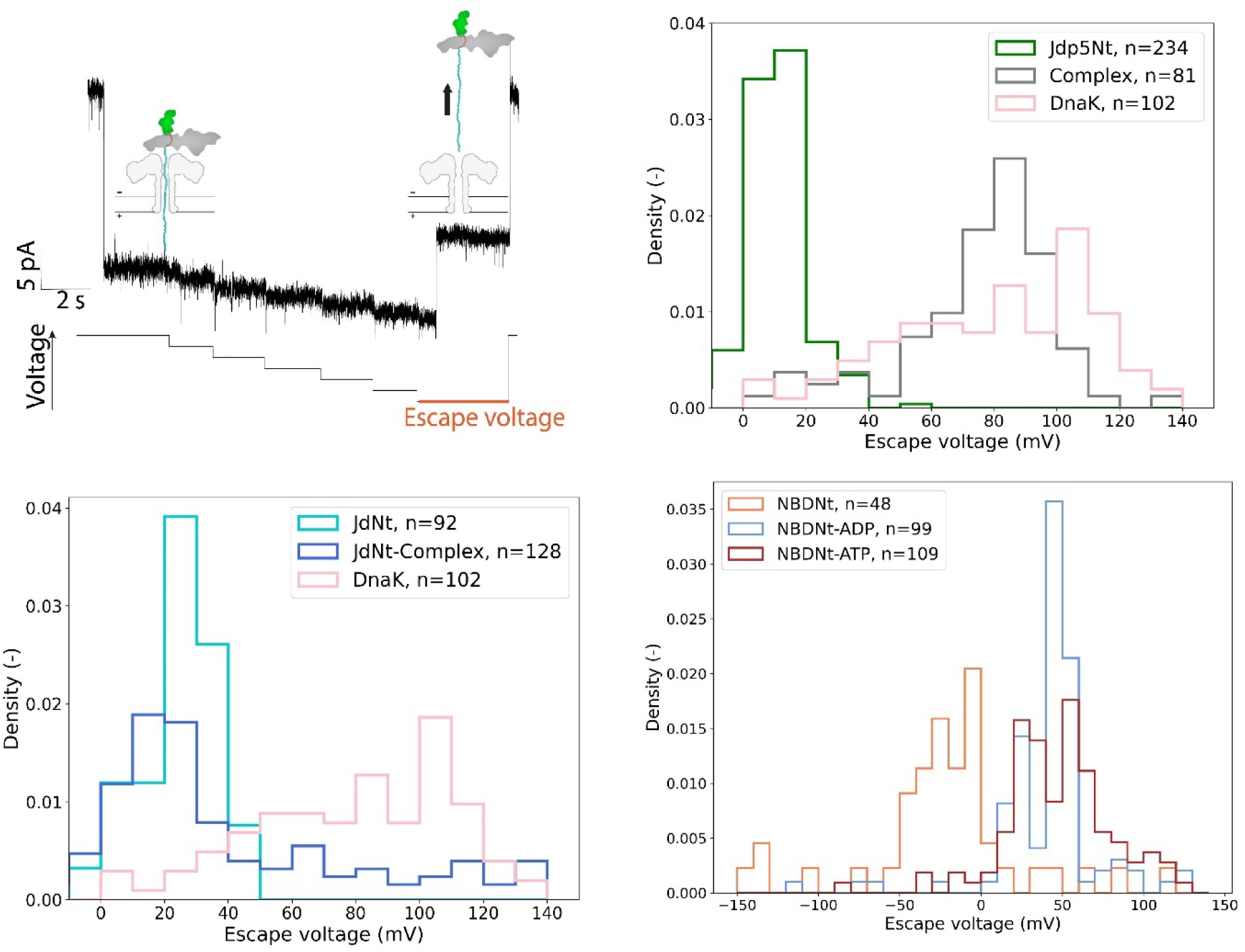
Escape voltage analysis. (a) illustration of escape voltage measurements. Ions can freely travel the aerolysin nanopore in its open state. Upon capture of an analyte (here the DnaK-JdptNt complex) a reduction in the measured current values is observed. Subsequently the voltage is lowered in 10 mV increments every 2 seconds until an upward shift in current values indicates the escape of the analyte from the pore. The voltage that was applied while the analyte escaped is the escape voltage of the trapping event. Histograms of the distribution of escape voltages are shown for measurements in the presence of Jdp5Nt, mixture of Jdp5Nt, DnaK, MgCl_2_ and ATP as well as DnaK on its own (b) and JdNt, mixture of JdNt, DnaK, MgCl_2_ and ATP as well as DnaK on its own (c). In (d) distributions of Escape voltages for NBDNt in its apo-state as well as in the presence of ADP or ATP are shown.

**Fig. S6.**
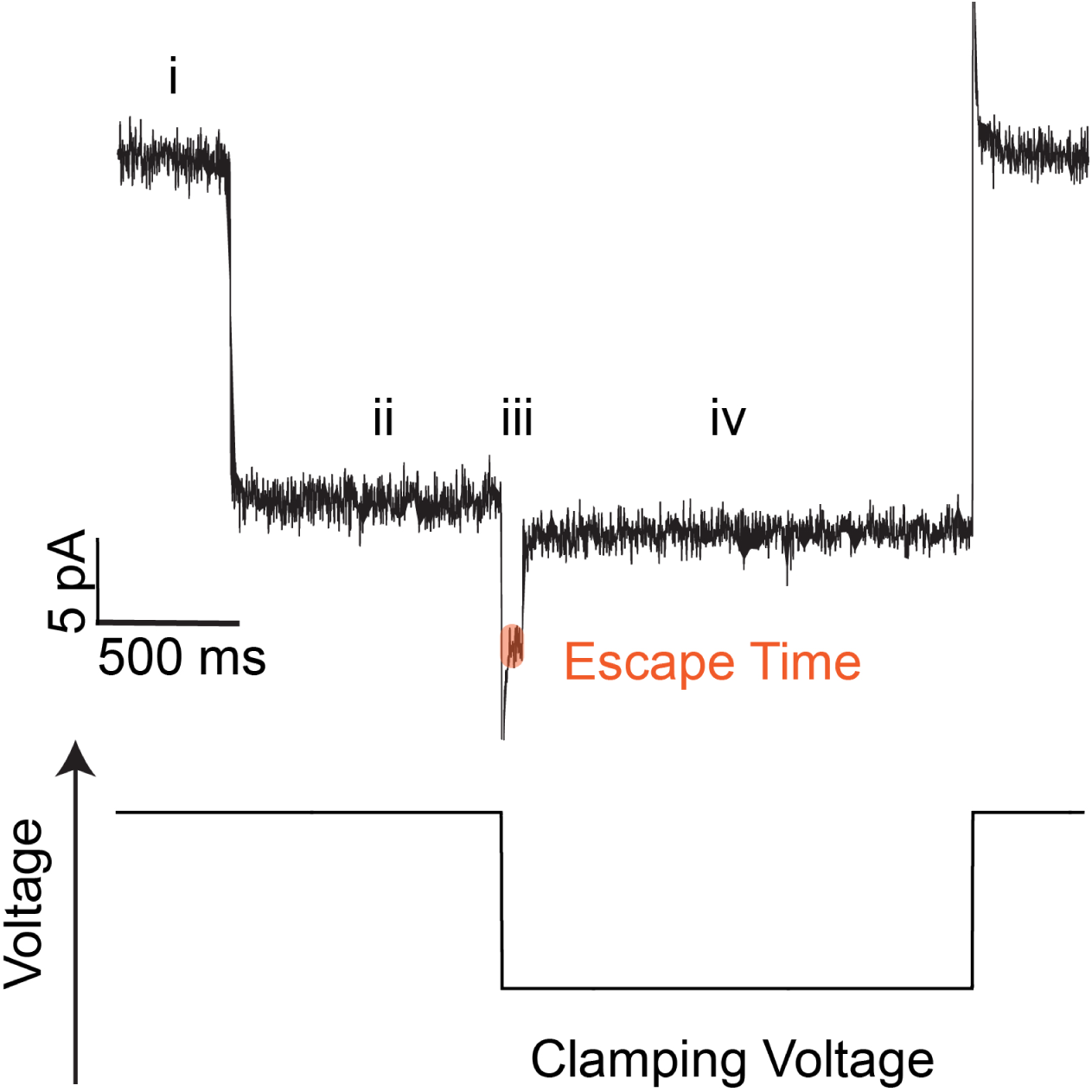
An example of escape time measurement for an event of Jdp5Nt-DnaK-complex trapping. An instantaneous reduction of the current signal from the baseline (i) to the analyte-specific blockage current (ii) is observed upon trapping. The voltage is subsequently lowered to the clamping voltage and the escape time (highlighted in orange) between the voltage switch and the escape of the analyte. When the escape occurs, the current is again instantaneously changed from the trapped current (iii) to the current of the unblocked pore at the clamping voltage (iv).

**Fig. S7.**
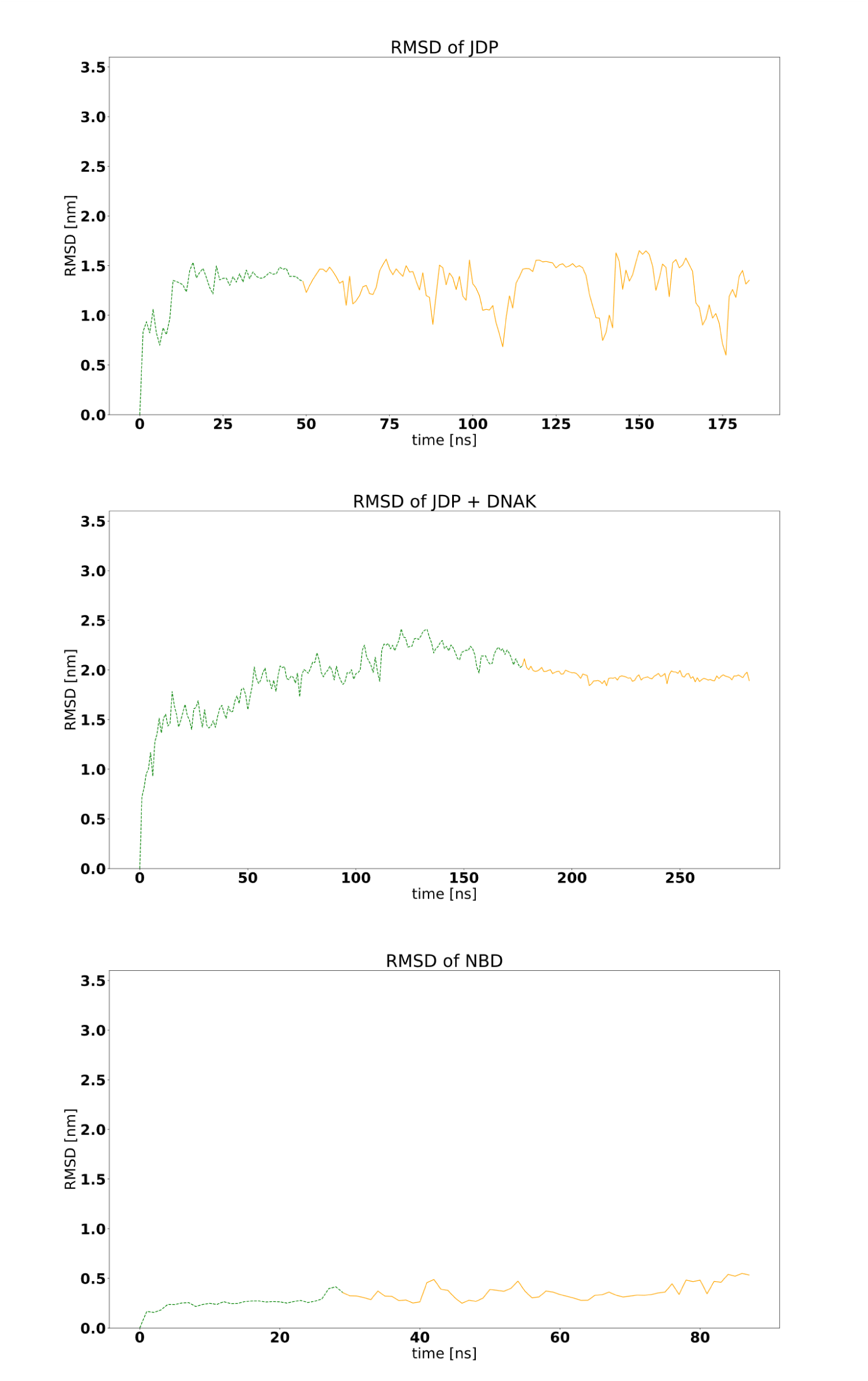
RMSD of each simulated system. The part of the trajectory used for averaging is indicated in yellow.

**Fig. S8.**
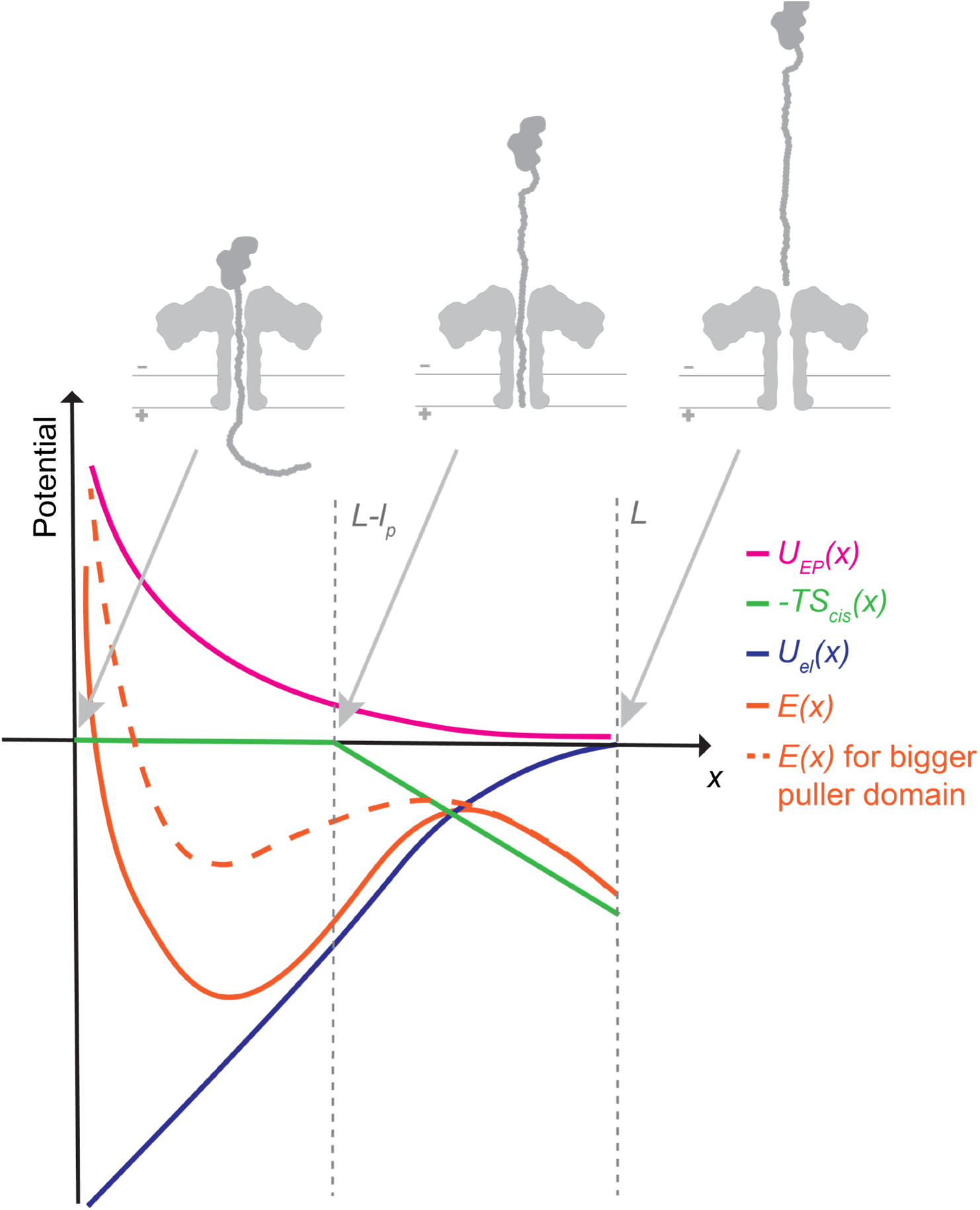
Illustration of the theoretic model of the energy landscape. With electrostatic potential 𝑼_𝒆_(𝒙), entropic pulling contribution 𝑼_𝑬_(𝒙) and the contribution of the unstructured threading tail = −𝑻𝑺_𝒄_(𝒙).

**Fig. S9.**
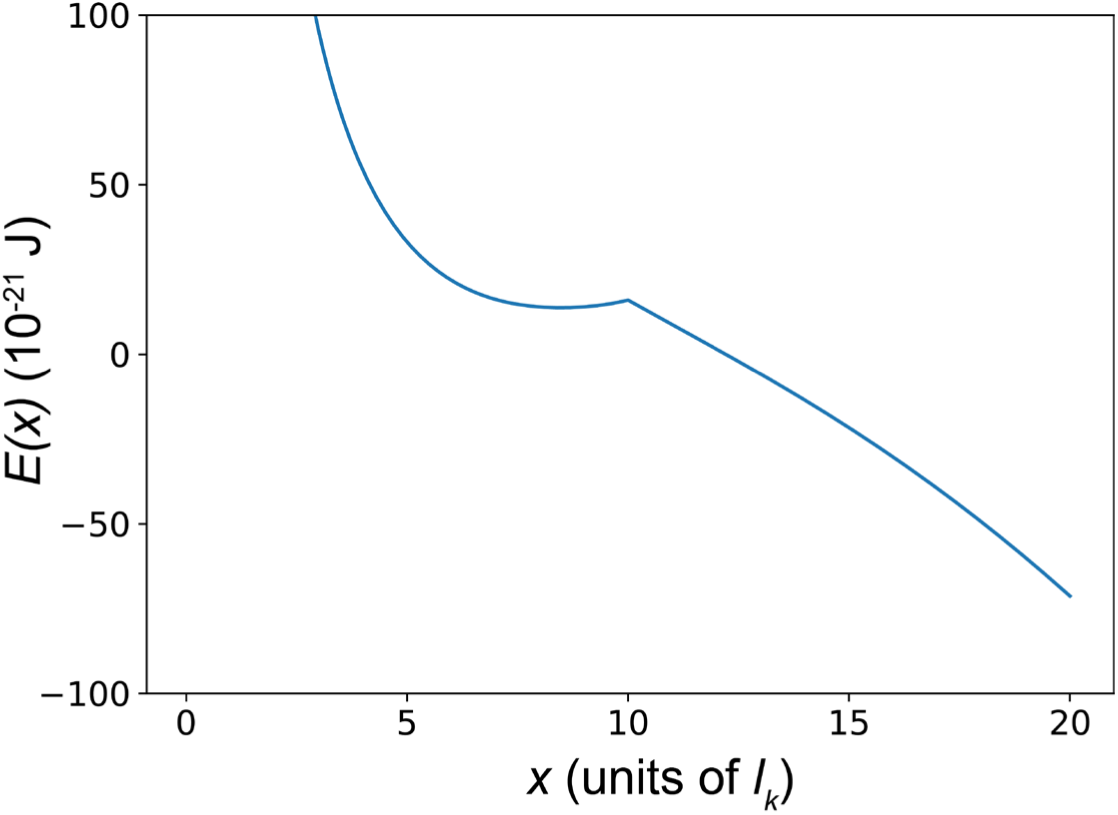
Energy landscape. 𝑬(𝒙) **for a puller with R = 10 *l_K_* at 55 mV clamping voltage.** With *x*, the length of the polypeptide that emerges from the pore on the *cis* side in units of the Kuhn length *l_K_* of an unstructured polypeptide.

**Table S1.**
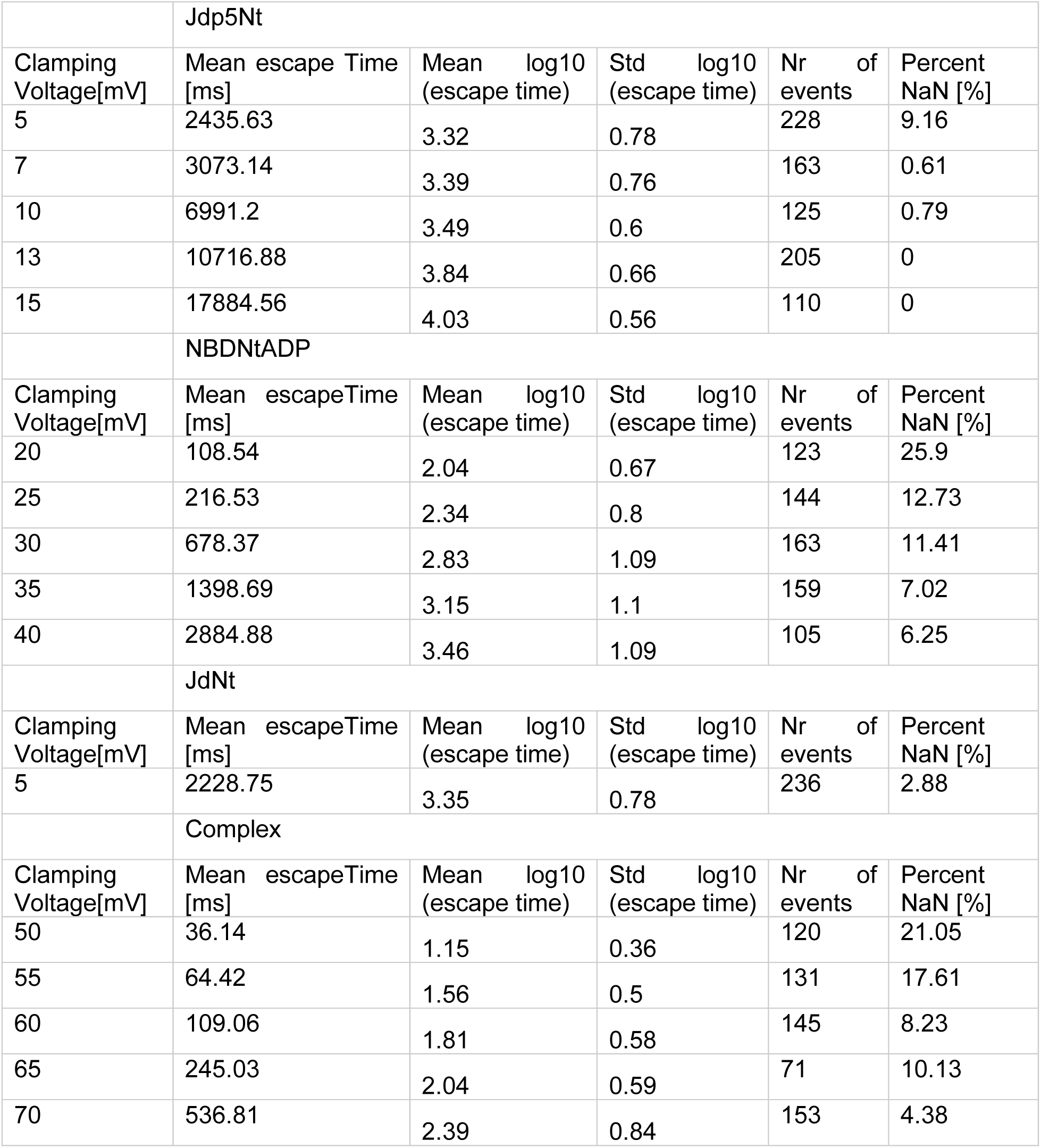
Mean escape values converted to ms, as well as mean and standard deviations calculated from log values. The number of values that were averaged for each analyte and clamping voltage are shown. Percentage NaN refers to the percentage of recorded trapping events, for which the escape was too fast to be detected.

**Table S2.**
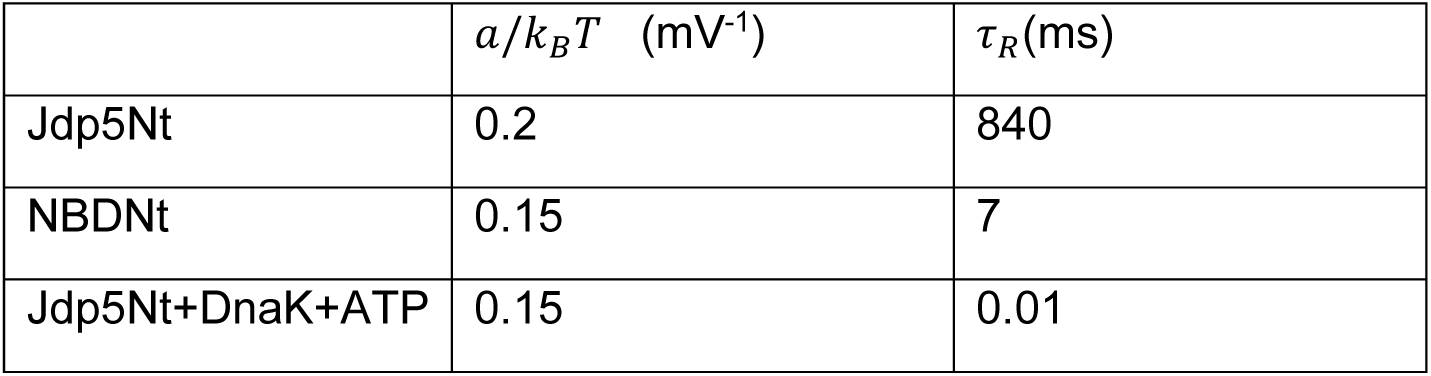
Extracted values from fitting experimental average escape times to Eq.2.

**Table S1.**
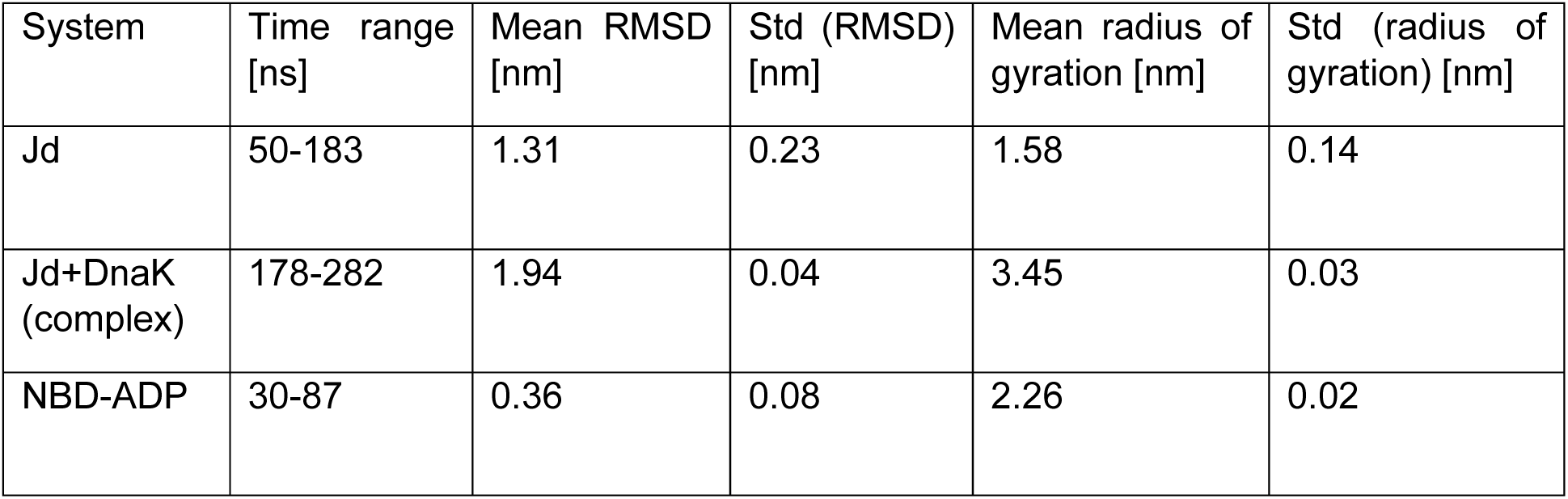
Summary of molecular dynamics simulations. Time ranges of trajectory used for radius, as well as average RMSD and radius of gyration with standard deviations for each system.

