## Supplemental information file for "Single-molecule evidence of Entropic Pulling by Hsp70 chaperones"

**The PDF file includes:**

Material and methods

Supplementary text

Note S1 Measurements of escape voltage

Note S2 Simulations of the escape process

Figures S1-S9

Tables S1-S3

References

### Materials and Methods

#### Protein production

The polypeptide moves in an energy landscape $E(x)$ that is the sum of three contributions (Fig.S8):

$$E(x) = -TS_{cis}(x) + U_{el}(x) + U_{EP}(x)$$

1) The length of the polypeptide in the *cis* side contributes by its entropy, which increases with the lenth as

$$S_{cis}(x) = k_B \frac{x}{l_K} \ln(4\pi) \qquad\qquad x \geq L - l_P$$

Here we have assumed that the polypeptide can be considered as a Freely Jointed Chain composed of a number of segments which is given by the length of the polypeptide emerging on the *cis* side, $x$, divided by the Kuhn length. We have also assumed that each segment can freely rotate, and that thus its entropy is just $k_B \ln(4\pi)$. This contribution applies if the polypeptide emerges only from the *cis* side. As soon as it emerges also on the *trans* side, the amount of length lost on one side is gained on the other and the two effects mutually cancel out.

2) Considering the polypeptide to be uniformly charged with charge density $\lambda$, in the presence of an applied voltage $V$ the electrostatic potential $U(x)$ is:

$$U_{el}(x) = \begin{cases} \dfrac{\lambda}{2}\dfrac{V}{l_P}(L-x)^2 & L - l_P \leq x \leq L \\ \dfrac{\lambda}{2} V l_P + \lambda V (L - l_P - x) & 0 \leq x < L - l_P \end{cases}$$

3) The entropic pulling contribution $U_{EP}(x)$ is (*12*):

$$U_{EP}(x) = \frac{3}{2} k_B T \frac{R^2}{l_K x}$$

where $R$ is the radius of the bulky, N-terminal domain (or of the bound DnaK).

The overall parameters we have used to compute the potential were:

$L = 20\ l_K\ ,\ l_P = 10\ l_K\ ,\ \lambda = \frac{q}{l_K} = -1.6\ 10^{-19}\ \mathrm{C/nm}\ ,\ k_B T = 4\ \mathrm{pN\ nm}$

The energy $E(x)$ for $V = 55\ \mathrm{mV}$ and $R = 10\ l_K$ , as an example, is represented in Fig.S9.

The escape process was simulated as a random walk in the interval $0 \leq x \leq L$. The walker started at a position $x_0$ and in a time-step it could move from $x$ of an amount $\Delta x$ to the left with probability

$$p_{left} = \frac{1}{2} \min\left(1, e^{-(E(x-\Delta x)-E(x))/k_B T}\right)$$

and to the right with the probability

$$p_{right} = \frac{1}{2} \min\left(1, e^{-(E(x+\Delta x)-E(x))/k_B T}\right)$$

and would not move with probability $p_{not\ move} = 1 - p_{left} - p_{right}$ .

The random walk started from an initial random position $x_0$ that was chosen by running the simulation with a high trapping voltage (140 mV) for 10’000 time-steps. The applied voltage was then lowered to the desired value, and the simulation was run until $x > L$, and the escape time, namely the number of time-steps (in arbitrary units) was recorded. The average escape time was estimated over 100 runs of the simulation ($x_0$ was re-generated for each run) for each applied voltage and each value of the radius $R$.

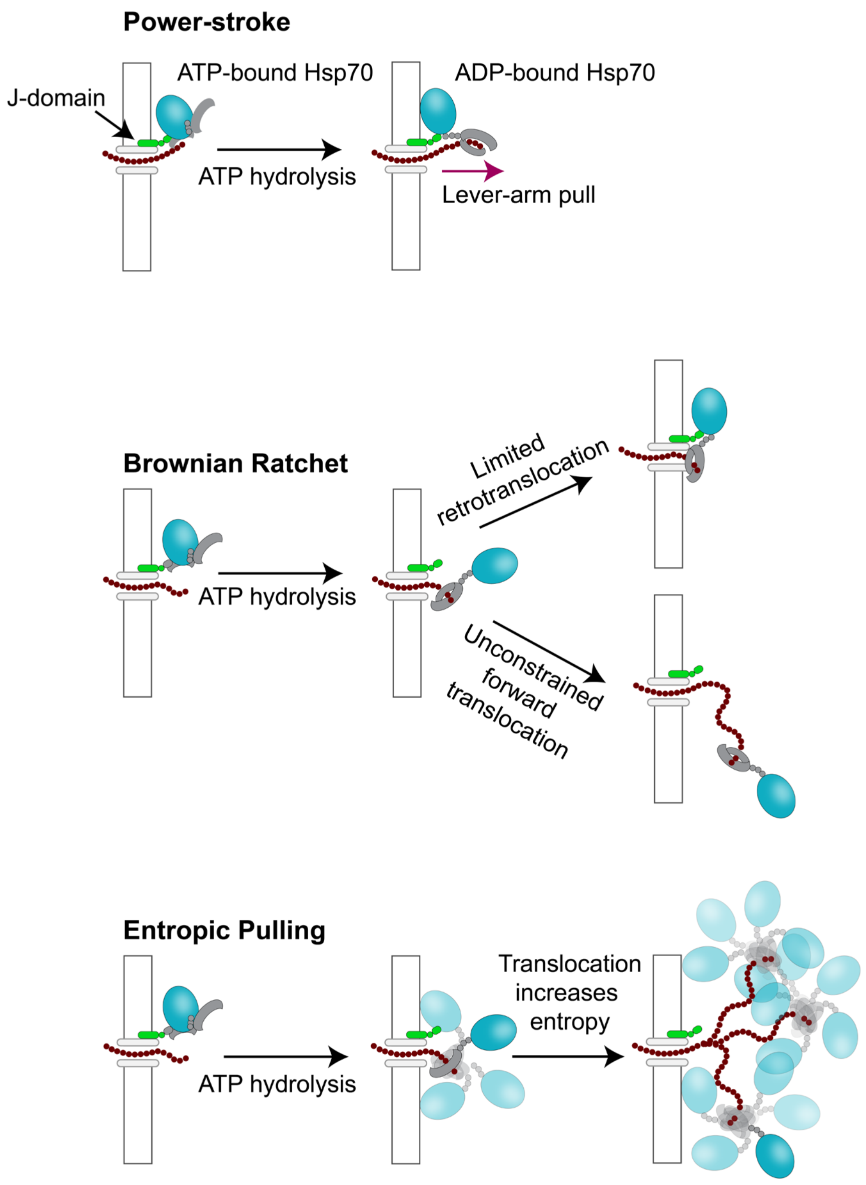


**Fig. S1 Proposed models for Hsp70s mechanism of function.**

Sequence of Jdp5Nt:

MKHHHHHHHHHHGSANAKQDYYEILGVSKTAEEREIRKAYKRLAMKYHPDRNQGDKEAEAKFKEIKEAYEVLTDSQKRAAYDQ
YGHAAFEQGG**ALLLSAPR**GSDGSESGDGGDSSEGSDGSESGDGGDSSEGSDGSESGDGGDSSEGSDGSESGDGGDSSE

Sequence of JdNt:

MKHHHHHHHHHHGSANAKQDYYEILGVSKTAEEREIRKAYKRLAMKYHPDRNQGDKEAEAKFKEIKEAYEVLTDSQKRAAYDQ
YGHAAFEQGGGSDGSESGDGGDSSEGSDGSESGDGGDSSEGSDGSESGDGGDSSEGSDGSESGDGGDSSE

Sequence of DnaK:

MGKIIGIDLGTTNSCVAIMDGTTPRVLENAEGDRTTPSIIAYTQDGETLVGQPAKRQAVTNPQNTLFAIKRLIGRRFQDEEVQRDV
SIMPFKIIAADNGDAWVEVKGQKMAPPQISAEVLKKMKKTAEDYLGEPVTEAVITVPAYFNDAQRQATKDAGRIAGLEVKRIINEP
TAAALAYGLDKGTGNRTIAVYDLGGGTFDISIIEIDEVDGEKTFEVLATNGDTHLGGEDFDSRLINYLVEEFKKDQGIDLRNDPLAM
QRLKEAAEKAKIELSSAQQTDVNLPYITADATGPKHMNIKVTRAKLESLVEDLVNRSIEPLKVALQDAGLSVSDIDDVILVGGQTR
MPMVQKKVAEFFGKEPRKDVNPDEAVAIGAAVQGGVLTGDVKDVLLLDVTPLSLGIETMGGVMTTLIAKNTTIPTKHSQVFSTAE
DNQSAVTIHVLQGERKRAADNKSLGQFNLDGINPAPRGMPQIEVTFDIDADGILHVSAKDKNSGKEQKITIKASSGLNEDEIQKMV
RDAEANAEADRKFEELVQTRNQGDHLLHSTRKQVEEAGDKLPADDKTAIESALTALETALKGEDKAAIEAKMQELAQVSQKLME
IAQQQHAQQQTAGADASANNAKDDDVVDAEFEEVKDKK

Sequence of NBDNt:

MKHHHHHHHHHHGSANMGKIIGIDLGTTNSCVAIMDGTTPRVLENAEGDRTTPSIIAYTQDGETLVGQPAKRQAVTNPQNTLFAI
KRLIGRRFQDEEVQRDVSIMPFKIIAADNGDAWVEVKGQKMAPPQISAEVLKKMKKTAEDYLGEPVTEAVITVPAYFNDAQRQAT
KDAGRIAGLEVKRIINEPTAAALAYGLDKGTGNRTIAVYDLGGGTFDISIIEIDEVDGEKTFEVLATNGDTHLGGEDFDSRLINYLVE
EFKKDQGIDLRNDPLAMQRLKEAAEKAKIELSSAQQTDVNLPYITADATGPKHMNIKVTRAKLESLVEDLVNRSIEPLKVALQDAG
LSVSDIDDVILVGGQTRMPMVQKKVAEFFGKEPRKDVNPDEAVAIGAAVQGGVLTGDVKDVLLLDGSDGSESGDGGDSSEGSD
GSESGDGGDSSEGSDGSESGDGGDSSEGSDGSESGDGGDSSE

**Fig. S2 Amino acid sequences of the proteins used in experiments.**

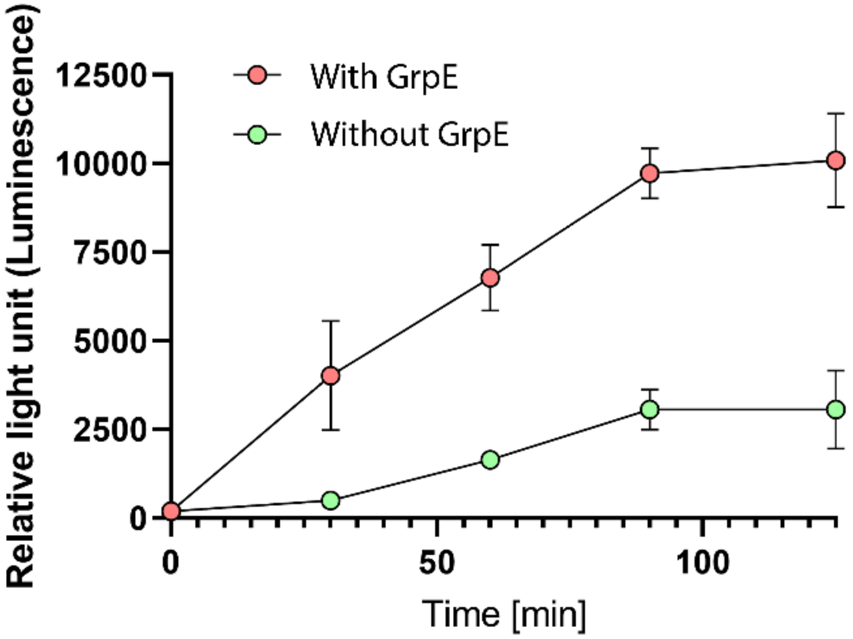


**Fig. S3 Time-dependent reactivation of urea-pre-denatured Luciferase that was incubated with (red) or without (green) GrpE.** 50 μM Luciferase was first inactivated for 2 min in 6M Urea, then diluted 125 folds in buffer (25 mM HEPES-KOH pH 7.5, 150 mM KCl, 10 mM MgAc2, 1mM DTT, and 5 mM ATP) at 30°C without any chaperone for 30 minutes. The reaction mix was then diluted 2 folds (final Luciferase concentration: 200 nM) and supplemented with the fully active chaperone mix (4 μM DnaK, 1 μM DnaJ with or without GrpE), to reactivate the aggregated luciferase at 25 °C. The error bars represent the SD between replicates (n = 3).

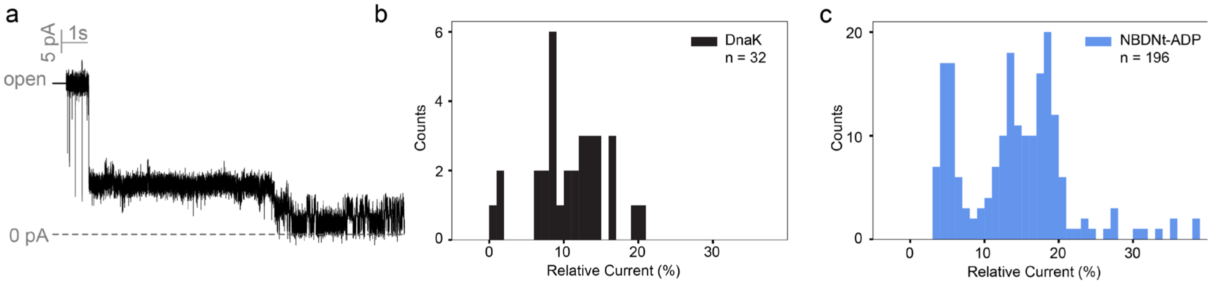


**Fig. S4 Current traces were acquired in 300mM KCl at 140 mV potential applied to the trans chamber filtered to 500 Hz.** (a): example trace recorded in a condition with Jdp5Nt, DnaK, $MgCl_2$ and ATP. After trapping a high current state associated with the Jdp5Nt-DnaK complex is observed, followed by the signal typical for Jdp5Nt. (b): Distribution of the fitted current values that were occasionally observed after addition of only DnaK. (c): Distribution of the fitted current values stemming from NBDNt in the presence of 1 mM ADP 10 mM $MgCl_2$.

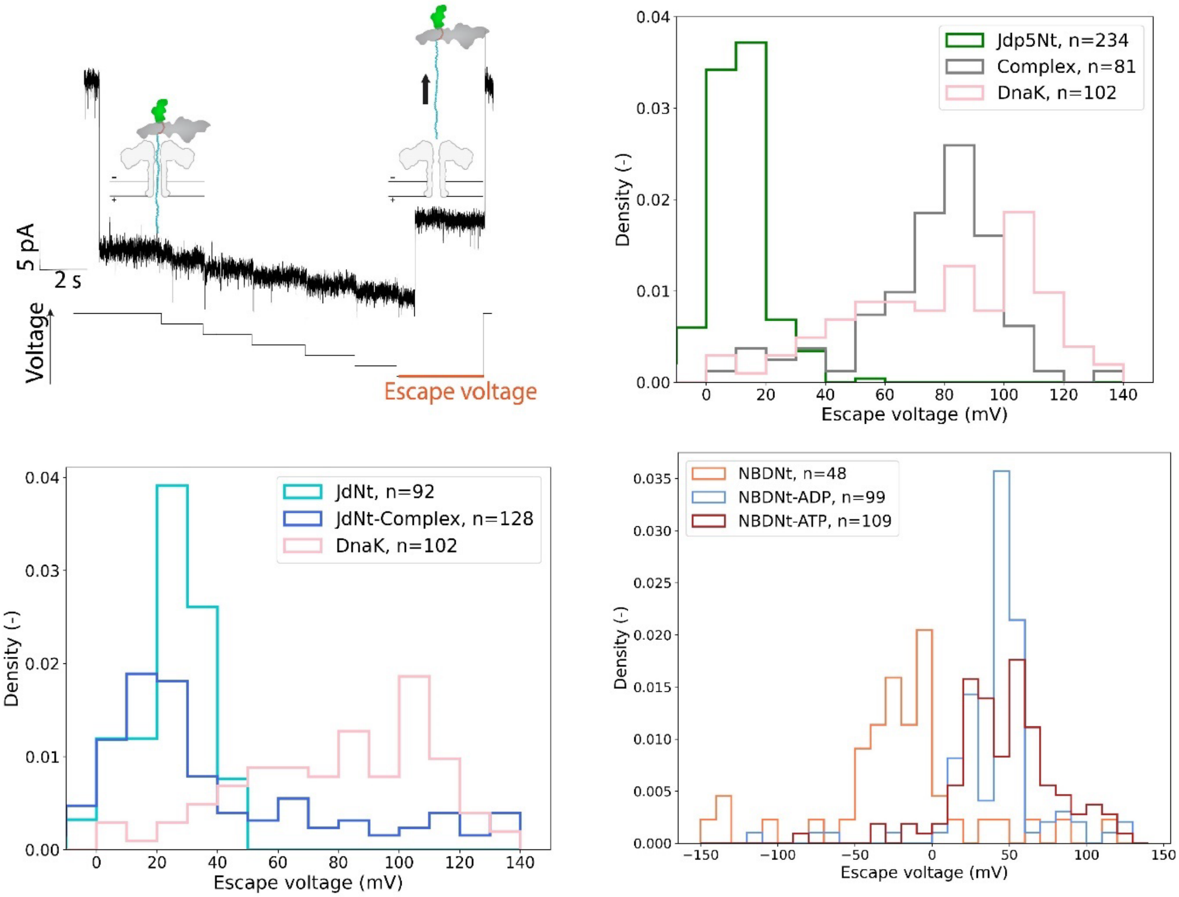


**Fig. S5 Escape voltage analysis.** (a) illustration of escape voltage measurements. Ions can freely travel the aerolysin nanopore in its open state. Upon capture of an analyte (here the DnaK-JdptNt complex) a reduction in the measured current values is observed. Subsequently the voltage is lowered in 10 mV increments every 2 seconds until an upward shift in current values indicates the escape of the analyte from the pore. The voltage that was applied while the analyte escaped is the escape voltage of the trapping event. Histograms of the distribution of escape voltages are shown for measurements in the presence of Jdp5Nt, mixture of Jdp5Nt, DnaK, $MgCl_2$ and ATP as well as DnaK on its own (b) and JdNt, mixture of JdNt, DnaK, $MgCl_2$ and ATP as well as DnaK on its own (c). In (d) distributions of Escape voltages for NBDNt in its apo-state as well as in the presence of ADP or ATP are shown.

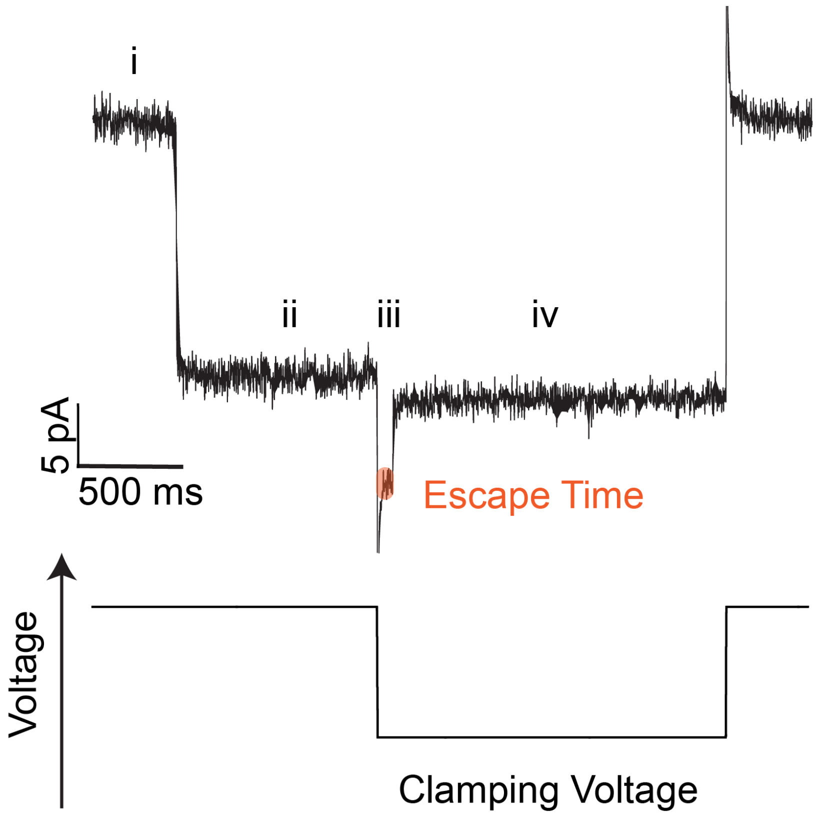


**Fig. S6 An example of escape time measurement for an event of Jdp5Nt-DnaK-complex trapping.** An instantaneous reduction of the current signal from the baseline (i) to the analyte-specific blockage current (ii) is observed upon trapping. The voltage is subsequently lowered to the clamping voltage and the escape time (highlighted in orange) between the voltage switch and the escape of the analyte. When the escape occurs, the current is again instantaneously changed from the trapped current (iii) to the current of the unblocked pore at the clamping voltage (iv).

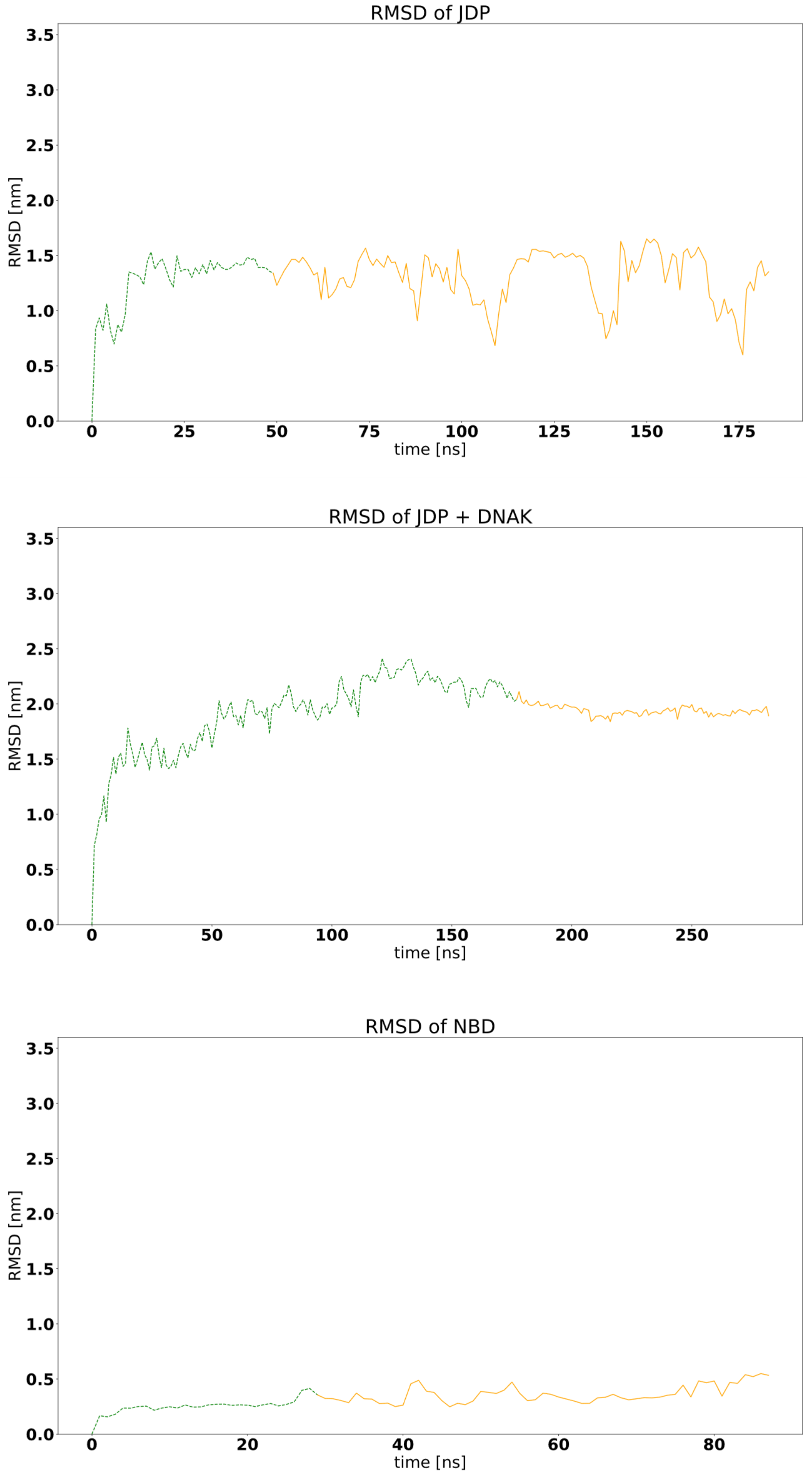


**Fig. S7 RMSD of each simulated system.** The part of the trajectory used for averaging is indicated in yellow.

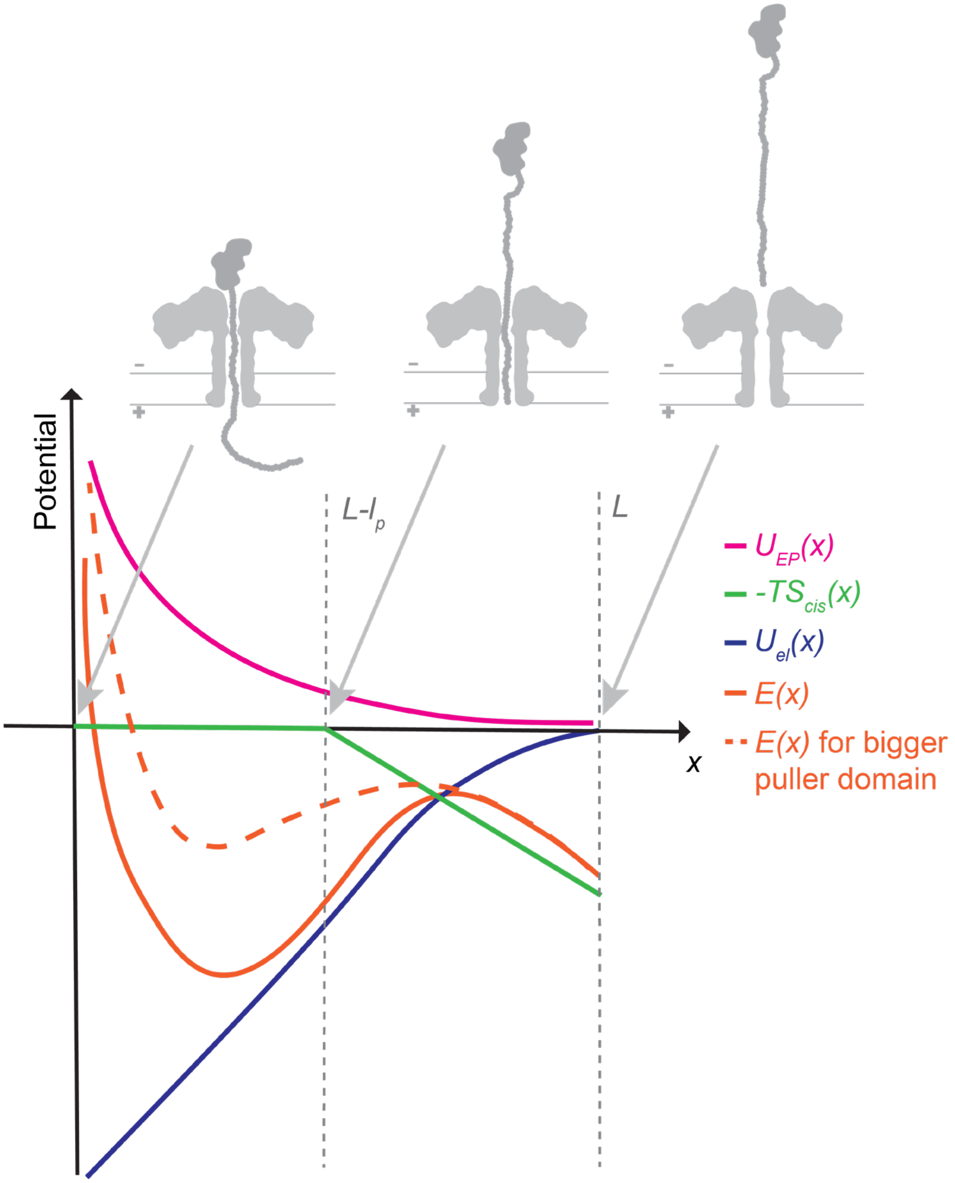


**Fig. S8 Illustration of the theoretic model of the energy landscape.** With electrostatic potential $\boldsymbol{U_{el}(x)}$, entropic pulling contribution $\boldsymbol{U_{EP}(x)}$ and the contribution of the unstructured threading tail $= \boldsymbol{-TS_{cis}(x)}$.

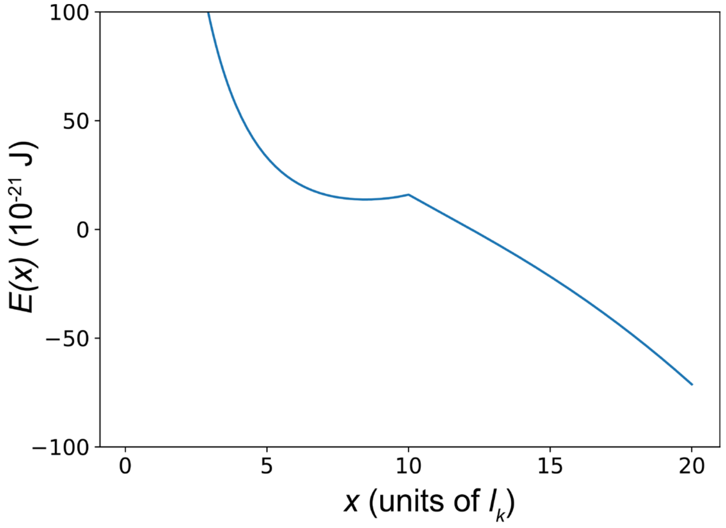


**Fig. S9 Energy landscape $E(x)$ for a puller with R = 10 $l_K$ at 55 mV clamping voltage.** With *x*, the length of the polypeptide that emerges from the pore on the *cis* side in units of the Kuhn length $l_K$ of an unstructured polypeptide.

**Table S1. Mean escape values converted to ms, as well as mean and standard deviations calculated from log values.** The number of values that were averaged for each analyte and clamping voltage are shown. Percentage NaN refers to the percentage of recorded trapping events, for which the escape was too fast to be detected.

| | Jdp5Nt | | | | |
|---|---|---|---|---|---|
| Clamping Voltage[mV] | Mean escape Time [ms] | Mean log10 (escape time) | Std log10 (escape time) | Nr of events | Percent NaN [%] |
| 5 | 2435.63 | 3.32 | 0.78 | 228 | 9.16 |
| 7 | 3073.14 | 3.39 | 0.76 | 163 | 0.61 |
| 10 | 6991.2 | 3.49 | 0.6 | 125 | 0.79 |
| 13 | 10716.88 | 3.84 | 0.66 | 205 | 0 |
| 15 | 17884.56 | 4.03 | 0.56 | 110 | 0 |
| | NBDNtADP | | | | |
| Clamping Voltage[mV] | Mean escapeTime [ms] | Mean log10 (escape time) | Std log10 (escape time) | Nr of events | Percent NaN [%] |
| 20 | 108.54 | 2.04 | 0.67 | 123 | 25.9 |
| 25 | 216.53 | 2.34 | 0.8 | 144 | 12.73 |
| 30 | 678.37 | 2.83 | 1.09 | 163 | 11.41 |
| 35 | 1398.69 | 3.15 | 1.1 | 159 | 7.02 |
| 40 | 2884.88 | 3.46 | 1.09 | 105 | 6.25 |
| | JdNt | | | | |
| Clamping Voltage[mV] | Mean escapeTime [ms] | Mean log10 (escape time) | Std log10 (escape time) | Nr of events | Percent NaN [%] |
| 5 | 2228.75 | 3.35 | 0.78 | 236 | 2.88 |
| | Complex | | | | |
| Clamping Voltage[mV] | Mean escapeTime [ms] | Mean log10 (escape time) | Std log10 (escape time) | Nr of events | Percent NaN [%] |
| 50 | 36.14 | 1.15 | 0.36 | 120 | 21.05 |
| 55 | 64.42 | 1.56 | 0.5 | 131 | 17.61 |
| 60 | 109.06 | 1.81 | 0.58 | 145 | 8.23 |
| 65 | 245.03 | 2.04 | 0.59 | 71 | 10.13 |
| 70 | 536.81 | 2.39 | 0.84 | 153 | 4.38 |

**Table S2. Extracted values from fitting experimental average escape times to Eq.2.**

| | $a/k_BT$ (mV$^{-1}$) | $\tau_R$(ms) |
|---|---|---|
| Jdp5Nt | 0.2 | 840 |
| NBDNt | 0.15 | 7 |
| Jdp5Nt+DnaK+ATP | 0.15 | 0.01 |

**Table S1. Summary of molecular dynamics simulations.** Time ranges of trajectory used for radius, as well as average RMSD and radius of gyration with standard deviations for each system.

| System | Time range [ns] | Mean RMSD [nm] | Std (RMSD) [nm] | Mean radius of gyration [nm] | Std (radius of gyration) [nm] |
|---|---|---|---|---|---|
| Jd | 50-183 | 1.31 | 0.23 | 1.58 | 0.14 |
| Jd+DnaK (complex) | 178-282 | 1.94 | 0.04 | 3.45 | 0.03 |
| NBD-ADP | 30-87 | 0.36 | 0.08 | 2.26 | 0.02 |
